# GMMA can stabilize proteins across different functional constraints

**DOI:** 10.1101/2024.01.30.577994

**Authors:** Nicolas Daffern, Kristoffer E. Johansson, Zachary T. Baumer, Nicholas R. Robertson, Janty Woojuh, Matthew A. Bedewitz, Zoë Davis, Ian Wheeldon, Sean R. Cutler, Kresten Lindorff-Larsen, Timothy A. Whitehead

## Abstract

Stabilizing proteins without otherwise hampering their function is a central task in protein engineering and design. PYR1 is a plant hormone receptor that has been engineered to bind diverse small molecule ligands. We sought a set of generalized mutations that would provide stability without affecting functionality for PYR1 variants with diverse ligand-binding capabilities. To do this we used a global multi-mutant analysis (GMMA) approach, which can identify substitutions that have stabilizing effects and do not lower function. GMMA has the added benefit of finding substitutions that are stabilizing in different sequence contexts and we hypothesized that applying GMMA to PYR1 with different functionalities would identify this set of generalized mutations. Indeed, conducting FACS and deep sequencing of libraries for PYR1 variants with two different functionalities and applying a GMMA analysis identified 5 substitutions that, when inserted into four PYR1 variants that each bind a unique ligand, provided 2-6°C in increased thermal stability and no decrease in functionality.

## INTRODUCTION

Improving the stability of a protein without impacting its desired function is a fundamental objective for protein engineering and design. Functional proteins often have strong constraints on their sequence. While approximately 5% of randomly sampled mutations can improve protein stability, far fewer do so without disrupting function[1,2]. As the benefits of each stabilizing mutation is small, whereas even a single destabilizing mutation can be highly detrimental, the protein designer must ‘shoot the moon’ by getting nearly all chosen mutations correctly. Earlier efforts incorporated evolutionary information in the form of consensus mutations or substitutions sampled often in the evolutionary history of the protein family[3–5]. More recently, protein design approaches have made incredible advances in predictions of changes in protein stability[6–8]. Many algorithms, most notably PROSS[9,10], sample only from a set of substitutions commonly seen throughout evolution. Such approaches have been successful for stabilizing many classes of proteins, including enzymes, transporters, and binding proteins[11– 13]. Still, identifying positions not to mutate and sets of substitutions that do not disrupt function is often performed empirically or with heuristics that may not transfer across to other systems. Such heuristics, like evolutionary conservation, may fail when applying design to engineered proteins with unknown functional constraints.

We have recently engineered over twenty new genetically encoded biosensors using the chemically induced dimerization PYR1-HAB1 partner proteins[14]. In plants, PYR1 is a soluble, 27 kDa receptor which recognizes the hormone abscisic acid (ABA)[15]. Upon noncovalent binding of ABA, PYR1 undergoes a subtle yet important conformational change that results in potent recognition of the PP2C phosphatase HAB1[16]. Extensive genetic analyses have revealed many mutations on PYR1 which result in constitutive binding to HAB1[17]. These mutations occur at the HAB1 binding interface, but also at distal positions. Thus, PYR1 is quite sensitive to mutations which disrupt function, hampering protein engineering and design efforts. It is unknown whether the conformation-induced mechanism is identical between the native and the engineered biosensors.

In this work, we identified amino acid substitutions that stabilized several already-developed PYR1 sensors of different specificity. We used a combined experimental and computational global multi-mutant analysis (GMMA), which has previously been shown to identify multiple substitutions that together progressively enhance a protein’s stability [18,19]. The ability to identify substitutions that work together is a result of analyzing variants with multiple amino acid substitutions where the beneficial effect is observed in various slightly different backgrounds. We hypothesized that this reduced sensitivity towards the sequence background would also enable GMMA to identify substitutions that will enhance PYR1 across sensors of different specificity. Applying GMMA to data generated by yeast display of PYR1 combinatorial mutagenesis libraries, we investigated the same sets of substitutions in two different sensor backgrounds and found a common set of mutations between the two sensors. We show that when these substitutions are incorporated into diverse sensors, they can also stabilize these proteins while not impacting *in vivo* and *in vitro* function.

## RESULTS

### Engineered PYR1 biosensors are less stable than PYR1

It is well established that directed evolution of proteins for new functions often lead to accumulation of thermally destabilizing mutations[20,21]. Proteins which accrue enough destabilizing mutations no longer fold at relevant temperatures, limiting many potential mutational trajectories. Thus, we hypothesized that many engineered PYR1 biosensors are less stable than the parental, abscisic acid-binding PYR1^ABA^. To test this hypothesis, we produced three engineered PYR1 biosensors, along with the parental PYR1 protein, as PYR1-Maltose Binding Protein (MBP)-His_6_ fusion proteins. Proteins were expressed in *E. coli* and purified by nickel affinity chromatography. PYR1^MANDI^ binds the agrochemical fungicide mandipropamid[22], while PYR1^DIAZI^ and PYR1^PIRI^ recognize the organophosphates diazinon and pirimiphos methyl, respectively[14]. We assessed thermal stability using a thermal inactivation assay. In this assay, a PYR1 biosensor was incubated at a given temperature for 20 minutes and then immediately assessed for ability to inhibit the phosphatase activity of HAB1 under saturating ligand concentrations. All three engineered biosensors were less thermally stable than PYR1^ABA^ (**Supplemental Fig 1**), supporting the hypothesis that accruing mutations which confer new binding specificities also decreases thermal stability. Therefore, we sought mutations that would stabilize the PYR1^ABA^ parental construct without destroying the ability of the protein to be engineered to bind diverse ligands.

### Cross enhancement strategy

To achieve enhancement across different sensors of different functional constraints, we chose to use a global multi-mutant analysis (GMMA). Previous work has shown that GMMA can identify up to six amino acid substitutions that enhance stability progressively under functional constraints, presumably because the substitutional effects are insensitive to the particular sequence background in which they are inserted[19]. We hypothesized that this insensitivity would also work across different functional constraints like the engineered PYR1-based sensors (**Figure 1**). GMMA takes as input a screen of many variants, each carrying several amino acid substitutions, and infers an average effect per substitution under an additive model of variant effects. We had previously developed a yeast display screening assay for PYR1 [23]. We used this assay to screen combinatorial libraries containing combinations of the same set of predicted stabilizing and destabilizing amino acid substitutions (**Figure 1A**). Combinatorial libraries were generated in two sensor backgrounds, PYR1^ABA^ and PYR1^DIAZI^. As discussed in more detail below, we used GMMA to identify enhancing substitutions for each sensor background and the results were combined resulting in a set of substitutions predicted to enhance both sensors. The cross-enhancement strategy is demonstrated by introducing the five best substitutions (HOT5) in two additional sensors PYR1^MANDI^ and PYR1^PIRI^.

**Figure 1.**
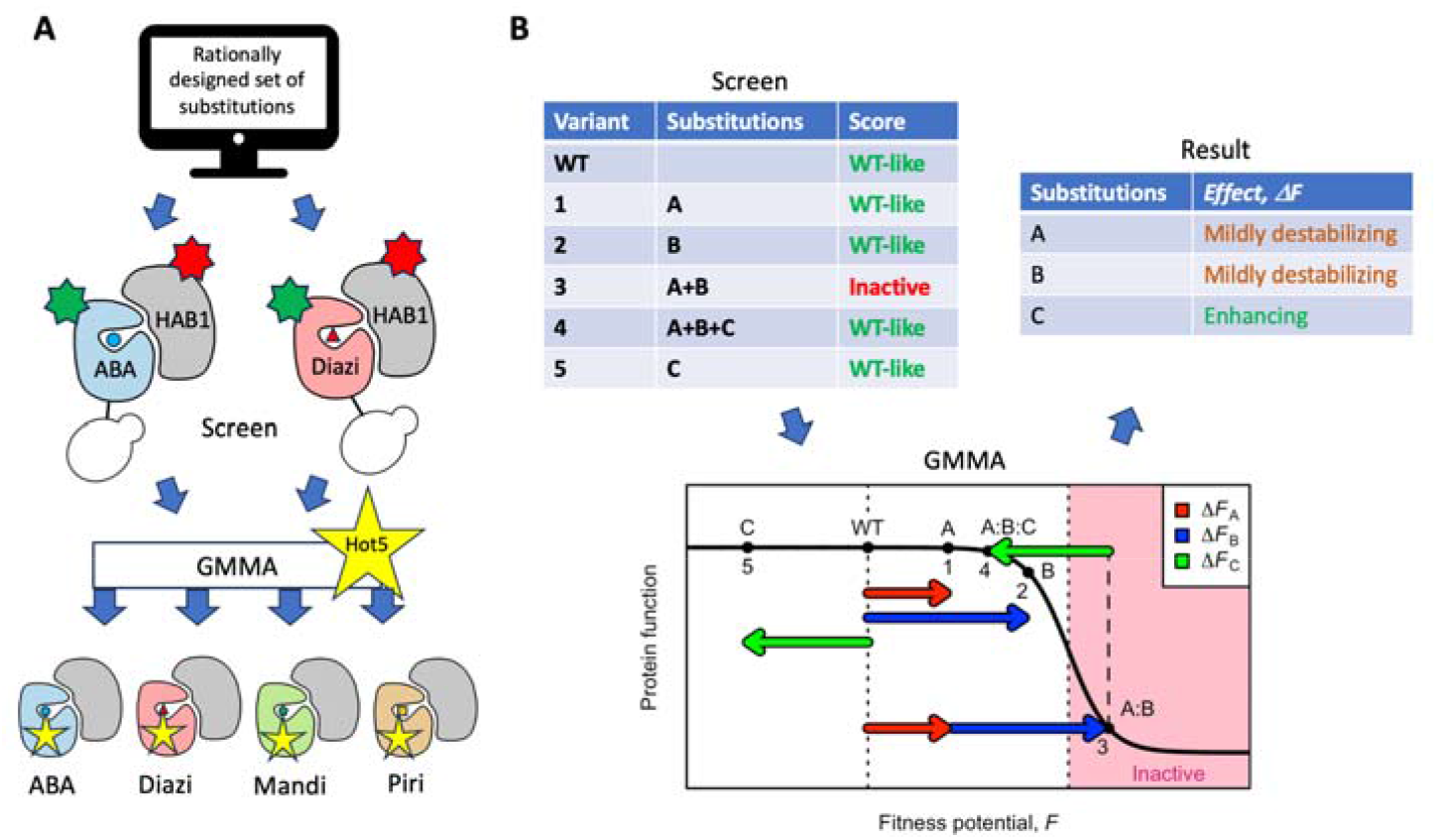
Cross enhancement strategy. (A) Cross-enhancement strategy: A single set of 99 amino acid substitutions is designed based on stability predictions, conservation, intuition, and guidelines for GMMA. The substitutions are combined randomly to make a large library of multi-mutants that is introduced into two sensor backgrounds and screened for ligand-mediated binding of HAB1 using yeast display, fluorescence activated cell sorting and deep sequencing. Based on the two screens, global multi-mutant analysis (GMMA) is used to identify a set of five amino acid substitutions for cross enhancement. These substitutions are introduced into a total of four sensor backgrounds for validation. (B) An illustration of GMMA involving five hypothetical protein variants (numbered 1–5) that are composed of mixtures of three amino acid substitutions (named A, B and C) each with a mild effect. In this example, all single-substitution variants are assayed to display wild-type like activity (variants 1, 2 and 5) from which beneficial substitutions are difficult to identify. By assuming additive effects (colored arrows) and a simple sigmoid global model (black line), the inactive double mutant A:B shows that both A and B are mildly destabilizing. Likewise, the substitution C may be inferred to be beneficial because it rescues activity in the triple-mutant A:B:C. A global fit seeks to estimate all effects, ΔF, such that the predicted functional level of variants match the screen. For more details and validation of GMMA see [18,19].

### Design of mutational libraries for GMMA

A central concept in GMMA is to identify enhancing substitutions by the ability to compensate for destabilizing ones. Thus, GMMA requires as input the functional screening of a library of multi-mutants containing a combination of predicted stabilizing and destabilizing mutations (**Figure 1B**). We sourced potential stabilizing substitutions identified from the Rosetta-based PROSS protocol from several solved PYR1 structures (see methods), from evolutionarily conserved positions, and by chemical and physical intuition. Our set of destabilizing mutations were predominantly large to small aliphatic substitutions in the protein core (see methods section). Additionally, we encoded reversion mutants for some of the ligand contacting positions in the engineered and less thermally stable PYR1^DIAZI^. In total, we assessed 99 substitutions at 74 of the 179 positions in the PYR1 encoding sequence (**Figure 2A, Supplemental Table 1**).

**Figure 2.**
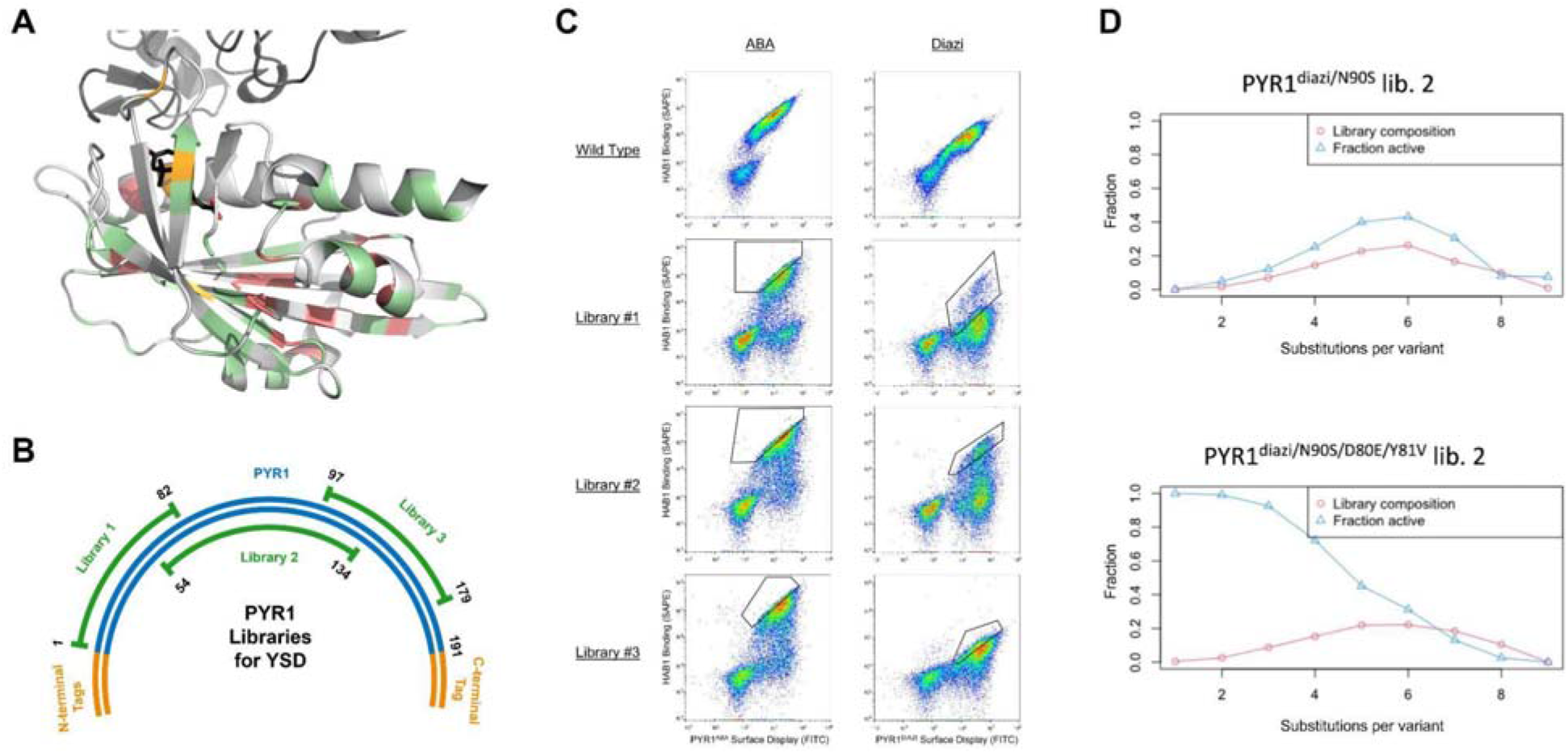
Design and screening of PYR1 mutagenic libraries. (A.) Positions of library substitutions on the structure of PYR1 (PDB entry 3QN1). The library consists of multiple substituted variants composed of 71 potentially stabilizing substitutions at 51 positions (green), 23 potentially destabilizing substitutions at 23 positions (red) and, for the PYR1^DIAZI^ background, 6 reversions (orange). Bound abscisic acid is shown as charcoal sticks. (B.) Library design strategy for screening. For each sensor, three separate combinatorial mutagenesis libraries were prepared. Each library covered a different gene section of PYR1. (C.) Binding cytograms for the six mutational libraries. The y-axis represents a fluorescence channel associated with phycoerythrin (PE) fluorophore (biotinylated HAB1^T+^, streptavidin PE conjugate), and x-axis represents the fluorescence channel associated with display of PYR1 as assessed by a C-terminal cmyc epitope tag and FITC conjugated anti-cmyc. Gates for the sorted population are shown in black. Conditions for screening were 500 nM of either abscisic acid (ABA) or diazinon (diazi) and 200 nM HAB1^T+^. (D) Multi-mutant composition of the Diazi sensor library 2 (red) and the fraction of binding variants (blue) as a function of the number of substitutions per variant. The same data is plotted using two different sequence references PYR1^DIAZI^ or PYR1^DIAZI/D80E/Y81V^.

The gene encoding PYR1 (537 bp) is too long for Illumina short read sequencing platforms. Therefore, we constructed three overlapping mutational tile libraries for PYR1^ABA^ and PYR1^DIAZI^ (**Figure 2B**). Tile library 1 covered positions 1-82, library 2 covered positions 54-134, and library 3 covered positions 97-179. We used combinatorial nicking mutagenesis[24,25] to generate library diversity, and bottlenecked each library to approximately 50,000 protein encoding variants to be able to sequence the libraries with reasonable coverage. We aimed for approximately a 50/50 ratio of functional and non-functional variants assumed to be optimal for GMMA.

Libraries were transformed into *S. cerevisiae* EBY100[26], protein induced with galactose, and biosensor variants screened using fluorescence activated cell sorting using previously optimized conditions for yeast display[23] (**Supplemental Fig 2**). The cytograms for libraries of PYR1^ABA^ and PYR1^DIAZI^ are shown in **Figure 2C** in comparison with the parental sensor backgrounds. The binding profiles for the parental sensors were as expected, with the yeast cells displaying the sensor showing a uniform HAB1^T+^ binding population that is ligand-dependent. In contrast, the mutational libraries showed ligand-dependent binding for only a subset of the displaying population. Many variants were non-functional as required for GMMA. Notably, the maximal binding signal in the PYR1^ABA^ and PYR1^DIAZI^ library 3 was much lower than the maximal binding signal in the other libraries. These results suggest that one or more substitutions common in libraries 1 and 2 between positions 54-82 increased the overall signal.

### Screen

For each library, we sorted half a million yeast cells by drawing a diagonal gate on the displaying and binding populations (see **Figure 2C** for gates used). These libraries were regrown, amplicons prepared, deep sequenced, and an enrichment score was calculated for each variant (see methods section; processed data is found in **Supplemental Data 1**).

GMMA benefits from statistical averaging of many variants that contain the same substitution in slightly different backgrounds (**Figure 1B**). The most informative region is when a substitution causes a variant to cross the major inactivation transition [19] where the readout is particularly sensitive to substitutions and thus, it is important that libraries contain an appropriate number of both stabilizing and destabilizing substitutions to populate this region[19]. In relation to this, all screens of PYR1^ABA^ libraries show compositions that are appropriate for GMMA with a range of 1–9 substitutions per variant that covers the transition from fully active to inactive variants (**Supplemental Fig 3**).

Surprisingly, the analysis of the PYR1^DIAZI^ libraries shows that substantially mutated variants have a higher fraction of active variants which is suboptimal for GMMA (**Figure 2D** and **Supplemental Fig 4**). Manual inspection revealed that most of the active variants from library 2 contain Y81V which is a revertant from the PYR1^DIAZI^ background. Many variants also contain D80E, but because this always co-occurs with Y81V, its individual effect is difficult to decipher. Because a high fraction of the variants contains these two substitutions it is possible to re-reference libraries 1 and 2 to PYR1^DIAZI/D80E/Y81V^ (9.5% of 3412 and 35% of 4483 variants respectively). The composition based on this reference sequence is far better suited for GMMA (**Figure 2D** and **Supplemental Fig 4**). This is consistent with the flow cytograms that show a lower signal for PYR1^DIAZI^ library 3 that do not contain mutations at positions 80 and 81 (**Figure 2C**).

### Identification of globally stabilizing mutations using GMMA

We performed GMMA as previously described [18,19] with an adjusted threshold for filtering confident substitution effects (see methods section). Previous applications of GMMA have only analyzed a single screened library and in the following we will investigate strategies for combining libraries in GMMA both within a single protein and across the two systems.

Enrichment scores calculated from the screens of individual libraries are normalized to the background (reference) sequence and thus comparable per sensor, provided that a common inactive signal is well populated in each library. Libraries for the PYR1^ABA^ sensor show similar enrichment score distributions which is a good indication that these are comparable (**Supplemental Fig 5**). Thus, we pool the enrichment scores for the three ABA libraries and perform a single GMMA. A comparison of GMMA results for individual libraries and the combined library confirms that the effects are reproduced well (**Supplemental Fig 6A**). For both approaches, GMMA estimates 38 substitution effects with high confidence of which 14 are inferred to be stabilizing.

As discussed above, PYR1^DIAZI^ enrichment scores for library 2 could only be normalized to the PYR1^DIAZI/D80E/Y81V^ reference, library 3 only to the original PYR1^DIAZI^ reference, while library 1 could be normalized to both. The distributions of enrichment scores confirms that library 2 is dependent on the shifted reference, with the majority of scores being better than the original PYR1^DIAZI^ reference sequence (**Supplemental Fig 7**). Thus, the Diazi analysis needed to be split into two or more GMMA’s. For PYR1^DIAZI/D80E/Y81V^ libraries 1 and 2, we find 76 confident substitution effects for both the individual and combined analyses, and effects are reproduced well between approaches (**Supplemental Fig 6B**). Of these, 19 are inferred to be stabilizing. In the GMMA analysis PYR1^DIAZI^ library 3 alone, we find 37 confident effects of which 4 are stabilizing.

Using the same reference sequence for Diazi libraries 1 and 3 results in similar score distributions (**Supplemental Fig 7**). However, the libraries have no overlap (common substitutions) in the GMMA and can therefore not be used in a single analysis (**Supplemental Fig 8A**). Because of this decoupling, it is in principle difficult to know if the scale of substitutions effect from library 3 is the same as for library 1, although a strong correlation between effects from combined library 1+2 and libraries 1+3 indicates that scales are comparable (**Supplemental Fig 8B**). Comparison of GMMA effects for the two sensors indicates that these reproduce the zero-point (separating stabilizing and destabilizing substitutions) but may be on a different scale (**Supplemental Fig 9**) and thus, the following selection focuses on ranks and stabilization within uncertainty.

To identify substitutions for cross-enhancement, we look for effects that enhance both the ABA and Diazi sensors in the combined analysis (**Figure 3** and **Supplemental Table 2**) but mention the individual analyses where we find it relevant (**Supplemental Fig 10**). In total, the combined analysis estimated 81 confident effects (87% of expected substitutions), of which 31 substitutions at 26 positions were identified as beneficial for at least one sensor (**Figure 3B**). These include substitutions spanning the length of the protein from E4 to A179; in helices, loops, and strands; and mostly in solvent exposed positions.

**Figure 3.**
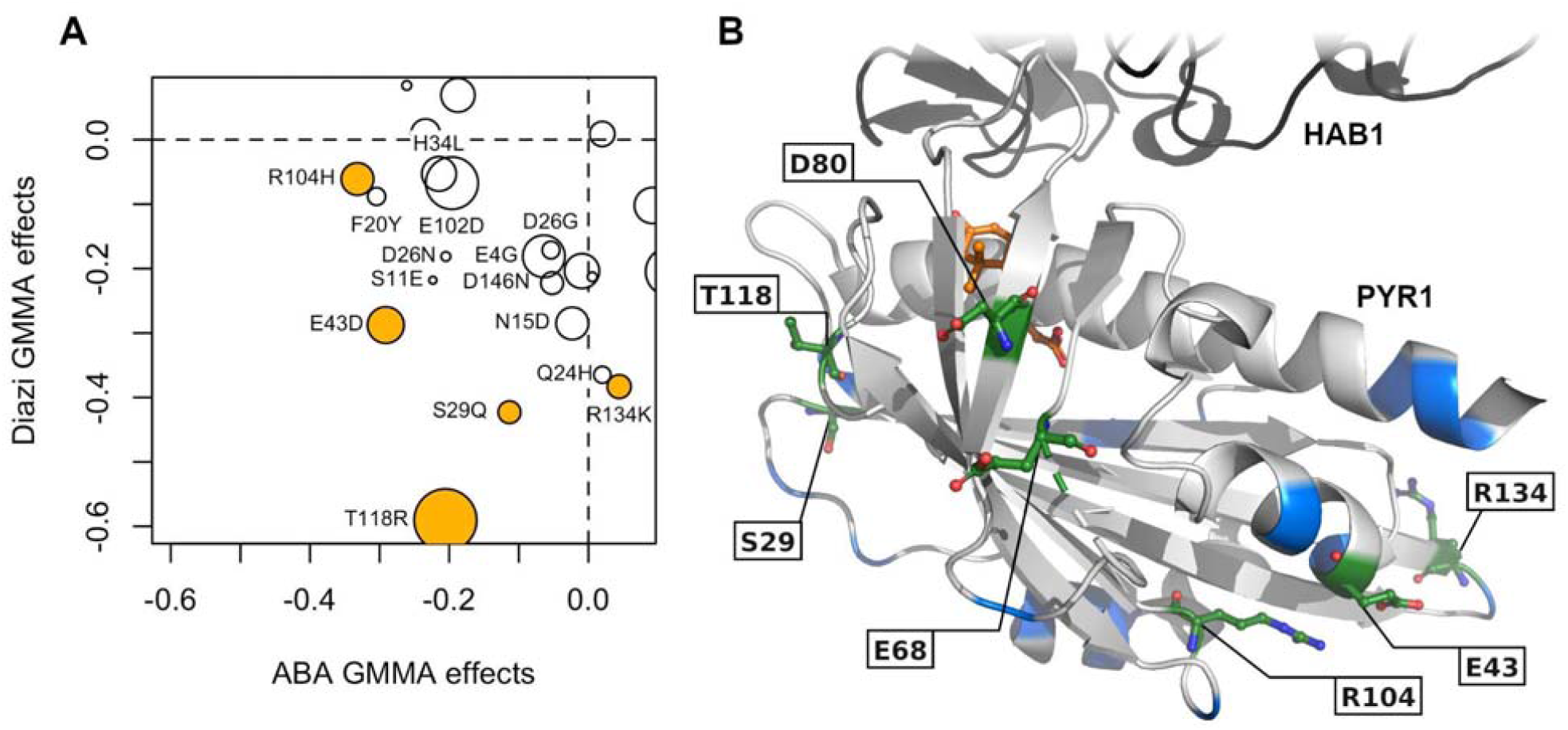
GMMA results combined for all libraries. (A) Point size scales inversely with a combined uncertainty and the selected substitutions are marked in yellow. The plot focuses on the stabilizing region of both sensors with many destabilizing substitutions not shown (all points shown in **Supplemental Fig 9**). Additional views were used in the final selection, see **Supplemental Fig 10** and **Supplemental Table 2**. (B) Structure of PYR1 (PDB entry 3QN1) showing the six selected substitutions (green). The ABA ligand is shown in orange and HAB1 in black. Other positions with stabilizing substitutions in one of the sensors are shown in blue.

The best substitution in the analysis of PYR1^ABA^, D80E, is contained in the optimized PYR1^DIAZI/D80E/Y81V^ background and thus identified as the best substitution for cross enhancement (together with the Diazi specific revertant Y81V). The only substitutions that are stabilizing within uncertainties in both sensors are T118R and E43D, with S29Q having slightly higher uncertainty in the ABA analysis. T118R is top ranking in three out of four individual analyses (**Supplemental Fig 10**) and has very low uncertainty in both individual and combined analyses (**Figure 3** and **Supplemental Table 2**). The two substitutions from library 1, S29Q and E43D, also show consistently good performance in both combined and individual analyses and are selected for cross enhancement. These four are perhaps the best substitutions and in the following we select two additional by less stringent criteria and note that this leaves several potentially well-performing substitutions untested (see discussion). Five of the six selected substitutions were predicted as stabilizing by PROSS; T118R was included based on evolutionary conservation.

Three substitutions from library 1, S11E, F20Y and D26N, are estimated to be enhancing in both sensors (**Figure 3A**), but the effects are typically exceeded by the uncertainty in both combined and individual analyses (**Supplemental Table 2** and **Supplemental Data 2**). Instead, we selected R104H and R134K that perform well in the individual analyses of library 2 and 3, respectively, for both sensors (**Supplemental Fig 10**). Furthermore, R104H ranks third in the combined analysis of the ABA sensor and R134K second in the Diazi sensor (**Supplemental Table 2**).

### GMMA designs are more stable than parental proteins

We ordered synthetic genes encoding eight designs in which we combined between two and four of the substitutions identified by GMMA and introduced these in the PYR1^ABA^ background.

Designs were expressed as genetic fusions with maltose binding protein. All expressed in soluble form and were purified in high yield. All designs were more stable than the wild-type PYR1 expression construct as judged by a thermal inactivation assay (Table 1). Three designs (2.1-2.3) contained two mutations from the set of S29Q, D80E, T118R. The designs (2.2, 2.3) containing the S29Q mutation were more stable than the design (2.1) that did not (Table 1). Four designs (3.1-3.4) shared D80E and T118R along with one of the four remaining mutations. Designs containing S29Q or R134K showed the highest thermal stability gain for the PYR1 sensor. We also tested a design (4) containing four GMMA-predicted stabilizing mutations which also provided stabilization.

**Table 1.** Stabilities of intermediate designs. Designs contained combinations of GMMA identified stabilizing mutations and were added to indicated PYR1 sensor backgrounds for testing. Data shown represents the average of 3 technical replicates.

| Design | Mutations | Stability change ($\Delta T_m$) - °C or yes/no | | | |
| --- | --- | --- | --- | --- | --- |
|  |  | PYR1<sup>ABA</sup> | PYR1<sup>DIAZINON/Y81V</sup> | PYR1<sup>MANDI</sup> | PYR1<sup>PIRI</sup> |
| 2.1 | D80E, T118R | 1.4 | NT | NT | NT |
| 2.2 | S29Q, D80E | 2.7 | NT | NT | NT |
| 2.3 | S29Q, T118R | 2.8 | NT | NT | NT |
| 3.1 | S29Q, D80E, T118R | 3.4 | NT | NT | NT |
| 3.2 | E43D, D80E, | 1.3 | NT | NT | NT |
|  | T118R |  |  |  |  |
| 3.3 | D80E, T118R,<br>R134K | 4.9 | Yes | Yes | Yes |
| 3.4 | D80E, R104H,<br>T118R | 1.0 | NT | NT | NT |
| 4 | S29Q, E43D,<br>D80E, T118R | 4.3 | Yes | Yes | Yes |

NT = not tested.

To examine the hypothesis that GMMA leads to the transferability of the stabilizing mutations, we tested the two most stable designs (3.3, 4) in the background of three engineered biosensors (PYR1^DIAZI^, PYR1^PIRI^, and PYR1^MANDI^). All six resulting proteins were more thermally stable than their parental sensor background (Table 1), demonstrating that GMMA-identified mutations are transferable across engineered proteins.

### HOT5 functions in vitro and generally stabilizes PYR1 background

We designed a set of substitutions, HOT5, combining five mutations (S29Q, E43D, D80E, T118R, R134K) from the two most stabilizing designs (3.3, 4). We tested the HOT5 mutations in four sensor backgrounds — PYR1^ABA^, the Y81V diazinon sensor revertant PYR1^DIAZI/Y81V^, PYR1^PIRI^, and PYR1^MANDI^. All parental backgrounds were included as controls. We assessed the ligand sensitivity using two different phosphatase inhibition assays (**Supplemental Fig 11**), PYR1-HAB1 inhibition in the absence of ligand, and the change in melting temperature as assessed by a thermal inactivation assay. A summary of the results is shown as **Figure 4**. The HOT5 background was functional in all four sensor backgrounds, did not change ligand sensitivity, and improved the stability of all proteins. The magnitude of the change was protein-dependent and ranged from 2°C (PYR1^MANDI^) to 6°C (PYR1^ABA^). Thus, the HOT5 background can stabilize engineered biosensors without negatively impacting function.

**Figure 4.**
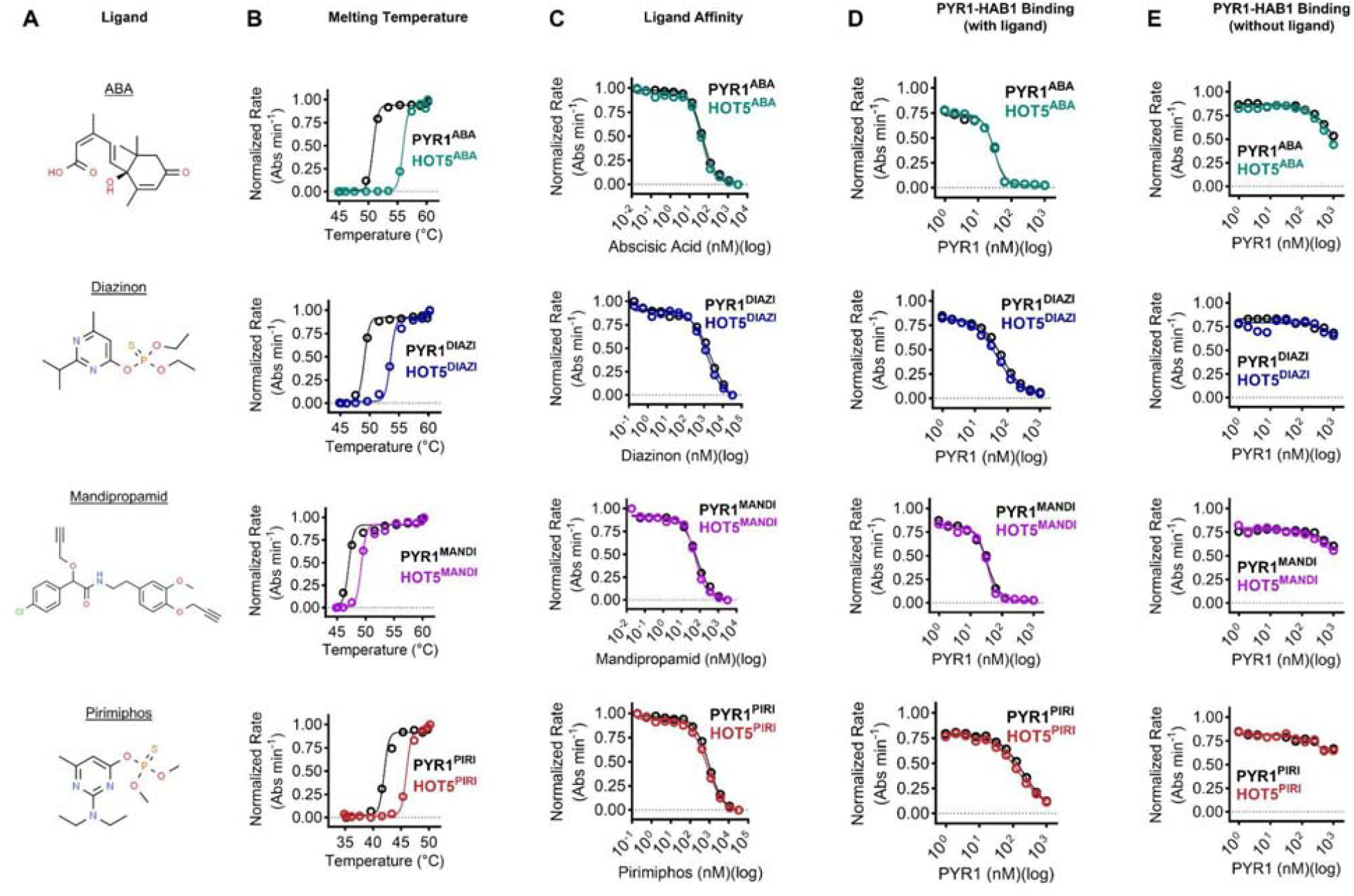
in vitro validation of HOT5 background for PYR1 biosensors. Each row shows the HOT5 sensor compared with its parental PYR1 sensor engineered to bind the indicated ligand. **A.** Ligand structures recognized by the respective sensor. **B.** Normalized HAB1 rate as a function of incubation temperature for four different sensor backgrounds. PYR1 sensors are incubated at the indicated temperature for 30 minutes and then removed and assessed for maintenance of activity by the ability to inhibit the phosphatase activity of HAB1 in the presence of respective ligand. **C.** Sensitivity of PYR1 sensors as assessed by ligand-dependent phosphatase inhibition assays. **D.** PYR1-HAB1 binding in the presence of 10 □M of respective ligand. **E.** Constitutive binding in the absence of ligand. HAB1 is fixed at 50 nM, with varying amounts of PYR1 protein indicated. For all panels, error bars are in all cases smaller than the symbols and represent 1 s.d., n=3 technical replicates.

To test whether HOT5 constructs can function *in vivo*, we developed a PYR1 genetic circuit in the yeast Kluyveromyces marxianus, which allows circuit testing at elevated temperatures. PYR1 is fused to a Z4 DNA-binding domain, while HAB1 is fused to the VP64 activation domain. Ligand-induced dimerization of PYR1 and HAB1 leads to transcription of the reporter eGFP. Compared with the original PYR1^DIAZI^ sensor, the HOT5 sensor allows a higher overall amount of gene expression at the highest diazinon concentrations at both 30°C and 42°C (**Figure 5A,** p< 0.01). For the pirimiphos sensors, both HOT5 and original versions yielded dose-dependent changes across the range of temperatures assessed, and the absolute magnitude of the response was similar between the two sensors (**Figure 5B**). Thus, HOT5 mutations can be transferred to different engineered biosensors and maintain function *in vivo* even at a variety of temperatures.

**Figure 5.**
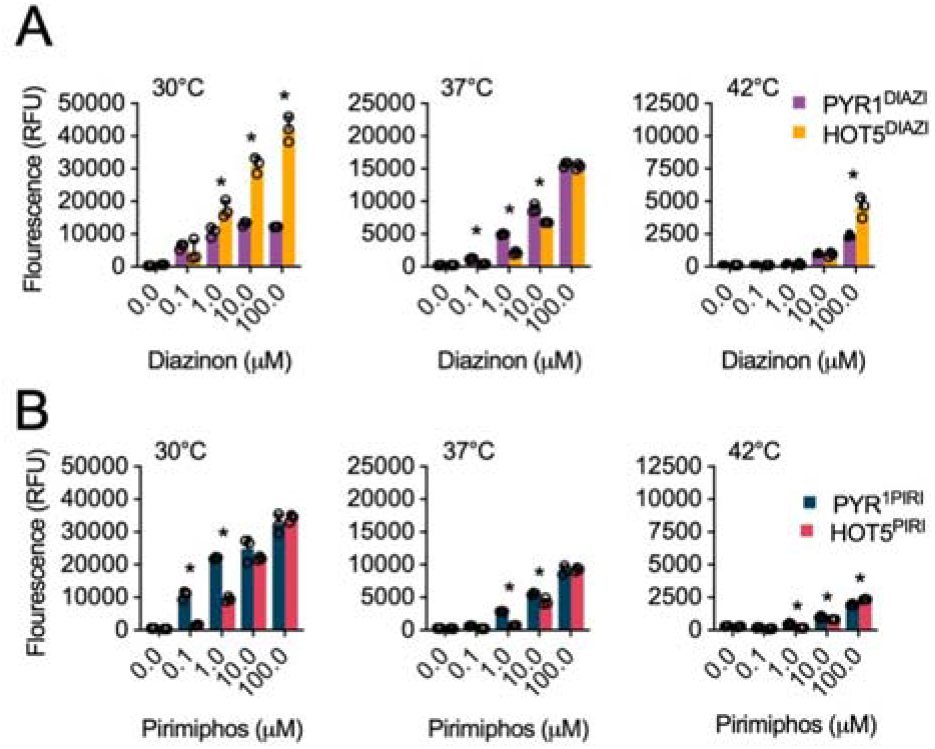
in vivo validation of Hot5 background for PYR1 biosensors. HOT5^DIAZI^ (A) and HOT5^PIRI^ (B) genetic circuit response in Kluyveromyces marxianus. (A) Z4-HOT5^DIAZI^ and VP64-HAB1 were expressed from a low copy number episomal plasmid in K. marxianus CBS6556 ABZ1::Z4_4_-EGFP URA3Δ HIS3Δ. Circuit activation was measured by quantifying EGFP expression. (B) HOT5^PIRI^ genetic circuit response in Kluyveromyces marxianus. Z4-HOT5^PIRI^ and VP64-HAB1 were expressed from a low copy number episomal plasmid in K. marxianus CBS6556. Circuit activation was measured by quantifying EGFP expression with Z4_4_-UAS-eGFP. For both panels, data points of biological triplicate are shown. * is p<0.01.

## DISCUSSION

A central challenge in the design of stable proteins is identifying sets of stabilizing mutations which do not otherwise impact function. The PYR1 protein is a prime example of this, as it undergoes subtle ligand-induced conformational changes essential for HAB1 recognition which can be disrupted by stabilizing mutations. We show that yeast display screening of combinatorial libraries, followed by GMMA analysis, is sufficient to recover sets of mutations which (i.) stabilize the PYR1 protein; and (ii.) maintain *in vitro* and *in vivo* function in a variety of engineered PYR1 backgrounds. We speculate that screening libraries of PYR1 variants harboring these mutations may rescue sensors for novel ligands.

The final design, HOT5, incorporated five substitutions from GMMA analysis. Four of these five came from the modified PROSS protocol, indicating the effectiveness of that protocol in identifying promising substitutions. Out of the 42 substitutions predicted by PROSS, 22 (52%) are non-destabilizing in both sensors compared to 28 out of all 65 (43%) substitutions predicted to be potentially stabilizing. These results confirm the power of PROSS to suggest putatively stabilizing variants that can then be refined further using methods such as GMMA.

The top-ranking substitution, D80E, consistently performs well in analyses of the ABA sensor; however, co-occurrence with Y81V in the Diazi libraries makes it difficult to separate the individual effects. All Diazi sensor variants that contain D80E also contain Y81V (but not vice versa). Notably, both individual and combined analyses of PYR1^DIAZI/D80E/Y81V^ show a large beneficial effect of the revertant E80D, though with a large uncertainty likely due to parameter co-variation in the global fit.

The upper stability limit for PYR1-based biosensors is most likely higher than the HOT5 design reported here. Many potentially stabilizing mutations were not incorporated in the combinatorial libraries used for screening. Additionally, several other candidates (E4G, S11E, F20Y and D26N) were not selected because their predicted effects have larger uncertainties, but may prove beneficial. Putting more emphasis on the initial analysis in GMMA[19], □*F*_*init*_, highlights two substitutions, E102D and D146N, that also perform well in the global analyses (**Supplemental Table 2** and **Figure 3A**).

This work demonstrates the power of GMMA analysis to identify substitutions that are stabilizing in engineered proteins like the PYR1-based biosensors. We speculate that similar GMMA approaches may find utility in identifying stabilized backgrounds for enzymes like cytochrome P450 oxygenases that have been engineered for non-native chemistries.

## MATERIALS AND METHODS

All plasmids, libraries, and primers used are listed in **Supporting Dataset 3**.

### Library construction

Potential stabilizing and destabilizing mutations were chosen from several sources. We employed PROSS with standard inputs and 6 distinct PYR1 structures from the PDB (PDB IDs: 3K3K, 3QN1, 4WVO, 5OR2, 3ZVU, 3K90). We looked for consensus mutations across the outputs. Mutations were then filtered by distance from ligand, distance from HAB1 interface, and lower contact number (more surface exposed). The PDB structure 3QN1 of the PYR1-HAB1 complex was analyzed for potentially stabilizing mutations using ChimeraX[27]. Mutations were chosen that increased internal hydrophobic packing, reduced buried charge, and added charged or polar residues to the protein’s exterior. Additionally, potentially stabilizing conserved PYR1 mutations were identified by performing sequence analysis of PYR1 homologs. Homologous sequences were identified using a BLAST search[28], aligned using Clustal Omega[29], and analyzed using Jalview[30]. Potentially destabilizing mutations were chosen by finding residues with less than 20% solvent-accessible surface area using the protSA web tool[31] and mutating them to residues with smaller, hydrophobic side chains (**Supplemental Table 1**).

Six separate mutational libraries were constructed using combinatorial nicking mutagenesis as described in Kirby et al[25]. Degenerate oligonucleotides encompassing one or more mutation sites were designed to contain both wild-type and mutant sequences and ordered from IDT. The resulting DNA was then transformed into chemically competent *S. cerevisiae* EBY100 cells[32] in SDCAA media and grown at 30 □. The cells were expanded to a 50 ml culture in fresh SDCAA media, grown to an OD_600_ of 1, transferred into yeast storage buffer (20% glycerol, 20 mM HEPES, 150 mM NaCl, pH 7.5), and stored at -80 □ as 1 ml aliquots.

### Yeast surface display and library sorting

To prepare libraries for sorting, cell aliquots were thawed on ice, spun down, and resuspended in SDCAA media to an OD_600_ of 1. Cells were then incubated at 30 □ for 4 hours before being spun down again, resuspended to an OD_600_ of 1 in 9 parts SGCAA and 1 part SDCAA media, and incubated overnight at 22 □. The next day, the cells were spun down, washed once with ice cold CBSF buffer (20 mM Trisodium Citrate, 150 mM NaCl, 5 mM KCl, 1 mg/ml BSA, pH 8), and resuspended to an OD_600_ of 2 in CBSF buffer. Cells were spun down again and stored as a pellet on ice until ready for labeling. Biotinylated HAB1 from ammonium sulfate stocks was spun down and resuspended to a concentration of 100 μM in CBSF containing freshly prepared 1mM DTT and 1 mM TCEP. Ligands were resuspended in ethanol (abscisic acid) or DMSO (diazinon) for sorting. Pelleted cells were resuspended to an OD_600_ of 2 in 1 ml of CBSF and incubated with 200 nM chemically biotinylated HAB1^T+^[14] and either 50 nM abscisic acid or 5 μM diazinon for 30 minutes at room temperature. Cells were then washed with 5 ml of CBSF, resuspended in 1.89 ml CBSF, and incubated on ice with 60 μl FITC and 50 μl SAPE on ice for 10 minutes. Cells were then washed with 5 ml of CBSF and stored as a pellet on ice. For sorting, cells were resuspended in 4 ml of CBSF, mixed by vortexing, and transferred to a 15 ml conical. 3 separate sorts were performed for each library with each sort gated for one of the following populations: all single cells, cells displaying PYR1, and cells bound to HAB1. Sorted cells were recovered in 5 ml of SDCAA media for approximately 45 hours at 30 □.

### Deep sequencing sample preparation

Library DNA from the sorted cells was prepped for deep sequencing exactly as described in Medina-Cucurella et al.[33]. Variants that were not observed in the unsorted pool were discarded and all pools were normalized to sequencing depth after a pseudo count of one was added. For variants with 20 or more reads (unnormalized sorted+unsorted), an enrichment score was calculated using:

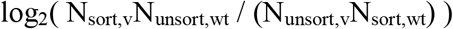

Since the scores are normalized to the same wild type (score zero) in all three libraries of the same sensor, we may compare scores across libraries if the depletion scale is similar in the libraries. In combined libraries, the average enrichment score is used if a variant is present in more libraries.

### Deep sequencing analysis

Deep sequencing reads were put through a quality filter and merged using a script written in-house, similarly as described in Haas et al.[34]. The number of reads containing each possible set of designed mutations were calculated and these numbers were used as inputs for the GMMA analysis.

## GMMA

GMMA was carried out as previously [18] with few settings adapted. The enrichment scores were used for GMMA to estimate effects of substitutions in the individual libraries and in combined libraries. Based on the enrichment score distributions, we fixed the activity levels in GMMA at zero (WT-like activity) and -7.0 (fully inactive) to make the global fits more robust (**Supplemental Fig 12**). The graph analysis of GMMA identified a set of three unexpected substitutions at positions 158-160 that was fully co-occurring and a few variants containing these needed to be removed to carry out the error analysis based on the Hessian matrix (3 and 15 variants from ABA and Diazi screens respectively). Similar to previously, we identified confident substitution effects as those observed in more than 20 variants[19]. Effects are reported as stabilizing if □*F* + *δ* < 0 and destabilizing if □*F* - *δ* > 0, where □*F* is the variant effect estimated by GMMA and *δ* is the uncertainty. For the combined ABA and Diazi analysis, a single uncertainty is calculated as the sum of variances. For the Diazi background, two GMMA’s were carried out; libraries 1 and 2 in PYR1^DIAZI/D80E/Y81V^ background and library 3 in unmodified PYR1^DIAZI^ background. Where substitution effects are estimated with confidence in both analyses (23 out of 80), the best is selected to represent that substitution in the combined analysis. In addition to the 99 substitutions that were expected, several unexpected substitutions were observed in the analyses, in total 265, 141 and 211 substitutions for ABA, Diazi tile 1+2 and Diazi tile 3 respectively. Many unexpected substitutions are rare and without confident estimates in GMMA, possible because these are in fact sequencing errors (**Supplemental Data 2**). Unexpected substitutions were never observed among the high ranking substitutions. Both individual and combined analyses resulted in 83 and 76 confident effects for ABA and Diazi tile 1+2 respectively.

### Construct creation for thermostable PYR1 sensors

Synthetic DNA (eBlocks; IDT) was ordered containing the coding sequences of all thermostable PYR1 variants and adaptor sequences that allowed for insertion into the pND004 MBP-tagged E. coli expression vector by Golden Gate assembly[35]. DNA was resuspended to a final concentration of 10 fmol/μl. 10 fmol DNA was then combined with 40 fmol of pND004 and added to a Golden Gate reaction mixture containing 20 units of BsaI-HF-v2, 400 units of T4 DNA ligase, and 1X T4 ligase buffer with a final total volume of 25 μl. The Golden Gate reaction was performed with 60 cycles alternating between 16 and 37 □ and two final incubation steps of 37 □ for 5 minutes and 65 □ for 10 minutes. 1 μl of the reaction mixture was pipetted directly into 20 μl of chemically competent Mach1 cells. The cells were then subjected to a scaled down version of a standard chemically competent *E. coli* transformation procedure using a 96-well plate for incubation steps and resuspending in 80 μl of LB for the recovery stage. Transformed cells were plated and incubated overnight at 37 °C, resulting in tens of colonies. The colonies were then grown up in 1 ml of LB and miniprepped.

### Protein purification

6xHis-MBP-ΔN-HAB1-C186S-C274S and 6xHis-MBP-ΔN-HAB1^T+^ were prepared exactly as described[23]. Thermostabilized PYR1 constructs were transformed into chemically competent BL21 (DE3) cells. Resulting colonies were used to inoculate 10 ml cultures and proteins were expressed by autoinduction as described previously[23] with an overnight 18 □ induction. The next day, the 10 ml cultures were centrifuged at 2500 x g at 4 □ in a swinging bucket rotor for 20 minutes. Prepared lysis buffer (100 mM HEPES, pH 8.0, 20% w/v Glycerol, 400 mM NaCl, 20 mM Imidazole, 30 mM MgCl_2_, 1.0 mg lysozyme per mL, 10 U TurboNuclease per mL, 1 μL 1 M PMSF per mL, 1 μL 1 M DTT per mL, 5 μL 1 M TCEP per mL) was mixed with 2X B-PER at an equivolume ratio immediately before use, and BPER-lysis buffer was added to cell pellets at 5 ml per 1 g of wet cell weight. Resuspended pellets were mixed in lysis buffer until there were no visible clumps and then transferred to a 50 ml Falcon tube and incubated at room temperature for 15 minutes. The lysed cells were then centrifuged at 20,000 x g for 20 minutes at 4 □. In the meantime, 0.5 ml of 50% slurry Ni-NTA resin per protein to be purified was placed in gravity flow columns and equilibrated with wash buffer (50 mM HEPES pH 8.0, 10% w/v Glycerol, 500 mM NaCl, 20 mM Imidazole, 1 mM DTT, 5 mM TCEP). The soluble supernatant from the spun down lysed cells was added to the columns and rocked on ice for 1 hour. The columns were then washed with 5 column volumes (2.5ml for 0.5ml resin) of wash buffer three times. The protein was then eluted from the column with 5 column volumes of elution buffer (50 mM HEPES pH 8.0, 10% w/v Glycerol, 200 mM NaCl, 500 mM Imidazole, 1 mM DTT, 5 mM TCEP). Samples from all steps of the purification were assessed by SDS-PAGE. Elutions were stored at 4 □ until use.

### Thermal melt and activity assays

Ligands were stored as 10 mM stocks in either ethanol (ABA) or DMSO (diazinon, mandipropamid, pirimiphos). Prior to assays, **buffer A** (20 mM Tris pH 7.9, 20 mM NaCl), **buffer B** (20 mM Tris pH 7.9, 20 mM NaCl, 2 mM MnCl_2_, 2 mM DTT, 0.6 mg/ml BSA), and **buffer C** (20 mM Tris pH 7.9, 20 mM NaCl, 2 mM MnCl_2_, 2 mM DTT, 0.6 mg/ml BSA, 10 mM p-nitrophenyl phosphate) were prepared and stored on ice. PYR1 proteins were exchanged into buffer A using PD-10 columns and their concentration was adjusted to 20 μM. Ammonium sulfate stocks of 6xHis-MBP-ΔN-HAB1-C186S-C274S were spun down, resuspended in buffer B to a concentration of 50 μM, and stored on ice.

For thermal melt assays, 40 μL PYR1 proteins were subjected to a thermal melt for 20 minutes at varying temperatures using a PCR machine. HAB1, ligands, and PYR1-post thermal melt were combined in wells of 96-well plates to a total volume of 100 μL and such that the final concentrations of all non-buffer components prior to data collection would be 4 μM ligand, 100 nM HAB1, 400 nM PYR1 for the ABA, Diazi, and Mandi sensors, and 1.2 μM PYR1 for the Piri sensors. Plates were incubated at room temperature on a shaker for 20 minutes and then 40 μL of the sample mixtures were added to 160 μL of buffer C. The samples were quickly mixed, bubbles removed, and the absorbance of the wells at 405 nm was measured every 20 seconds for 10 minutes using a plate reader.

For PYR1 titration assays, stocks were made of HAB1 (500 nM) with either ligands (20 μM) or an equivolume amount of ethanol or DMSO as a control. 35 μL of 20 μM PYR1 stocks were then combined with 35 μL of the HAB1 +/-ligand stocks and this mixture was added to the first well of a 96-well plate. 70 μL of the HAB1 +/-ligand stocks were then added to all wells of the appropriate rows of the 96-well plate in order to perform serial dilutions with 2-fold dilution steps. Plates were incubated at room temperature on a shaker for 20 minutes and then 40 μL of the sample mixtures were added to 160 μL of buffer C. The samples were quickly mixed, bubbles removed, and the absorbance of the wells at 405 nm was measured every 20 seconds for 10 minutes using a plate reader. The final concentration of HAB1 was 100 nM and of the ligands (for non-control samples) was 4 μM.

For ligand titration assays, stocks were made of HAB1 (250 nM) with PYR1 (500 nM for ABA sensor, 1.25 μM for diazinon and mandipropamid sensors, and 5 μM for pirimiphos sensors). 150 μL of PYR1 + HAB1 stocks were added to the first well of a 96-well plate and 100 μL were then added to the rest of rows of the in order to perform serial dilutions with 3-fold dilution steps. Ligands were then added to the appropriate first well (333 nM ABA, 33 μM diazinon and pirimiphos, and 3.3 μM mandipropamid) and dilutions were performed. Plates were incubated at room temperature on a shaker for 20 minutes and then 40 μL of the sample mixtures were added to 160 μL of buffer C. The samples were quickly mixed, bubbles removed, and the absorbance of the wells at 405 nm was measured every 20 seconds for 10 minutes using a plate reader.

After collection, all data was processed, fit, and graphed using custom R scripts or GraphPad Prism.

### Strains and media Kluyveromyces marxianus

CBS 6556 *ura3Δ his3Δ* was used as a starting strain for all experiments described in this work. All constructed strains are listed in **Supplemental Table 3**. Synthetic defined (SD) media was used for all plasmid-based expression experiments. The SD-uracil (SD-U) medium is defined as 6.7 g/L BD Difco™ Yeast Nitrogen Base without amino acids, 1.92 g/L Yeast Synthetic Drop-out Medium Supplements without uracil, and 20 g/L D-glucose. SD-H and SD-H-U are similarly defined but with 0.75 g/L of CSM-His and CSM-His-Ura, respectively. For all pathway refactoring experiments and 2-PE biosynthesis analysis, *K. marxianus* strains were cultivated in rich medium (YPD: 10 g/L Gibco™ Bacto™ Yeast Extract, 20 g/L Gibco™ Bacto™ Peptone, 20 g/L d-glucose). 20 g/L agar was added to make solid agar plates. All yeast cultures were conducted in 250 mL baffled shake flasks containing 25 mL of appropriate media. Culturing was conducted in an INFORS HT Multitron incubation shaker with temperature control set to 30, 37, and 42 °C as needed.

### *K. marxianus* sensor systems and transformations

A *K. marxianus* EGFP reporter strain (Ys2042) was created by integrating a reporter cassette (HTB1p-Z4_4_-HTB1_core_-EGFP-CYC1t) at the ABZ1 locus in CBS 6556 ura3Δ his3Δ, described in detail by[36]. In short, two plasmids were transformed into K. marxianus. One contains Cas9 and the gRNA targeting ABZ1; the other contains 700 bp up- and downstream homology arms surrounding the reporter cassette. Integration was confirmed with colony PCR and Sanger sequencing. Sensor plasmids were created by PCR amplifying relevant PYR1 sensors with sw420 and sw422 then HiFi Gibson assembled (NEB) into the pSW310 backbone digested with BamHI and NheI. The sensor plasmid (PGK1p-VP64_AD_-ΔN-HAB1-PGK1t and TEF1-Z4_DBD_-PYR1-TDH1t) containing a URA3 auxotrophic marker was then transformed with a chemical transformation protocol into the Ys2042 strain, also described by[36]. In brief, 2 mL of late exponential phase cells were washed twice with water. To the cell pellet, 10 uL 10 mg/mL ssDNA (Thermo salmon sperm DNA) and 800 ng of plasmid DNA were added. Fresh transformation buffer (500 uL of 40% w/w autoclave sterilized polyethylene glycol 3350 (PEG), 0.45-μm filtered 0.1 M lithium acetate (LiAc), 10 mM Tris–HCl (pH 7.5), 1 mM EDTA (1□×□TE buffer), and 10 mM DTT) was mixed thoroughly with the cells and DNA and left at room temperature for 15 minutes. Heat shock was performed at 47 °C for 5 minutes. The supernatant was removed, and the cells were plated at dilutions on SD-U plates. After 30 hours, colonies were picked, grown in SD-U liquid culture, and stored as glycerol stocks.

### Flow cytometry analysis of sensor response in *K. marxianus*

*K. marxianus* strains containing biosensor constructs were streaked on SD-U agar and then a single colony was inoculated into 2 mL SD-U media containing 2% glucose in 14 mL culture tubes (Corning, Tewksbury, MA, USA). After overnight growth at 30 °C shaking at 220 RPM, pre-culture OD_600_ values were measured and then used to inoculate to an OD_600_ value of 0.05 in fresh SD-U media in a 96 deep-well plate (USA Scientific, Orlando, FL, USA). A maximum of 5 μL of ligand solvated in DMSO was added at various concentrations to the inoculated wells. The plate was then sealed with AeraSeal film (Excel Scientific, Victorville, CA, USA) and grown at 30 °C, 37 °C, or 42 °C, 1,000 RPM, 80% relative humidity for 12 hours. Cells were centrifuged at 5700 g for 3 minutes and resuspended in water with a final dilution factor of ¼ for flow cytometer analysis. A BD accuri™ C6 flow cytometer (BD Bioscience) was used to collect and analyze data. 10,000 events were collected for each sample, and the forward scatter, side scatter, and EGFP fluorescence (530 nm bandpass filter) were recorded.

### Receptor and protein expression and purification

BL21 (DE3) cells expressing MBP fusions to either PYR1^WT^ HOT5^WT^, PYR1^DIAZI^, HOT5^DIAZI^, and PYR1^PIRI^, and HOT5^PIRI^ were individually cultured in 400 mL LB-kanamycin media to an OD_600_ ∼ 0.7 and then induced with 1 mM IPTG for 20 hours at 22°C. Cells were pelleted, resuspended in 15 mL of lysis buffer (50 mM HEPES pH 8.0, 200 mM NaCl, 10 mM Imidazole, 10% Glycerol), freeze-thawed three times, and then sonicated and cellular debris removed by centrifugation. The supernatants were loaded onto 0.5 mL Ni Superflow resin columns, washed three times with 15 mL of lysis buffer, and then eluted (50 mM HEPES pH 8.0, 500 mM imidazole, 200 mM NaCl, 10% Glycerol, 1 mM MnCl_2_). Fractions with an A_260_/A_280_ of less than ∼1 and a protein concentration greater than ∼1 mg/mL were pooled together and dialyzed overnight in 1 L of dialysis buffer (50 mM HEPES pH 8.0, 200 mM NaCl, 10% Glycerol, 0.1% 2-mercaptoethanol). Dialyzed protein solutions were diluted with glycerol to a final 40% glycerol composition and stored at -80°C. 6xHis-ΔN-HAB1 was expressed using a pET23 expression vector as previously described[23]. HAB1 Phosphatase expression and purification conditions were the same as receptor proteins with the exception that an MnCl_2_ cofactor (1 mM) was added to LB media, lysis buffer, elution buffer, and dialysis buffer.

### ΔN-HAB1 phosphatase inhibition assays

PYR1 and HOT5 proteins were titrated against ΔN-HAB1 to determine relative concentrations of active receptors to phosphatase before subsequent phosphatase inhibition assays. Protein titrations were conducted in a 96-well plate with ΔN-HAB1 (50 nM for PYR1^WT^, HOT5^WT^, HOT5^DIAZI^, 20 nM for HOT5^PIRI^), 4-methylumbelliferone phosphate substrate (1 mM), varying concentrations of MBP-tagged recombinant receptor proteins (0 nM to 200 nM, or 0 nM to 400 nM for HOT5^PIRI^), and their corresponding ligands (10 uM), ABA, diazinon, or pirimiphos methyl. Curves were fitted to a 4-parameter log-logistic model to determine EC_50_ values of each receptor, which were compared to HAB1 to find relative ratios of active receptor to phosphatase. Relative percentages of active HAB1 to receptors were 81% for PYR1^WT^, 52% for HOT5^ABA^, 105% for HOT5^DIAZI^, 776% for HOT5^PIRI^. Subsequent phosphatase inhibition assays used the corresponding concentrations of active proteins. Phosphatase inhibition assays were conducted with varying concentrations of chemical (ABA, diazinon, or pirimiphos methyl) ranging from 19.5 nM to 100,000 nM, followed by 10 nM ΔN-HAB1 in phosphatase assay buffer (100 mM Tris-HCl –pH7.9, 100 mM NaCl, 3 mg/ml BSA, 0.1% 2-mercaptoethanol, 1 mM MnCl_2_). Reactions were mixed and equilibrated for 5 minutes before the addition of substrate (1 mM 4-methylumbelliferone phosphate) and measurement. Fluorescence data were collected (λ_ex_ = 360 nm, λ_em_ = 460 nm) on a Tecan Spark multimode microplate reader.

## Acknowledgements

We thank Prof. Jakob R. Winther for discussions and insights related to GMMA. This work was supported by Defense Advanced Research Projects Agency Advanced Plant Technologies (DARPA APT, HR001118C0137) to T.A.W., S.R.C., and I.W; National Science Foundation GRFP to Z.T.B; the PRISM (Protein Interactions and Stability in Medicine and Genomics) centre funded by the Novo Nordisk Foundation (NNF18OC0033950) to K.L.L. K.E.J. and

K.L.L acknowledge access to computational resources at the Biocomputing Core Facility at the Department of Biology, University of Copenhagen. The views, opinions, and/or findings expressed are those of the authors and should not be interpreted as representing the official views or policies of the Department of Defense or the U.S. Government. Approved for Public Release, Distribution Unlimited.

## Author Contributions

Conceptualization: N.D., K.E.J., K.L.L., T.A.W.; Designed bench research: N.D., N.R.R., J.W., M.A.B., I.W., S.C., T.A.W.; Performed bench research: N.D., Z.T.B., N.R.R., J.W., M.A.B.; Developed computational algorithms: K.E.J., K.L.L.; Developed novel code: K.E.J.; Data analysis: N.D., K.E.J., Z.T.B., Z.D.; Writing: N.D., K.E.J., K.L.L, T.A.W. with input from all co-authors; Supervision: I.W., S.R.C., K.L.L., T.A.W.; Funding Acquisition: I.W., S.R.C, T.A.W, K.L.L..

## Data Availability

All scripts are available on GitHub https://github.com/KULL-Centre/_2024_Daffern_GMMA_PYR1. Processed deep sequencing data is available on the sequencing read archive. SRA accession number: PRJNA994968

