## Supplementary Materials. for "GMMA can stabilize proteins across different functional constraints"

**This document contains:**

**Supplemental Figures 1-13**
**Supplemental Tables 1-3**

### Supplemental Figures

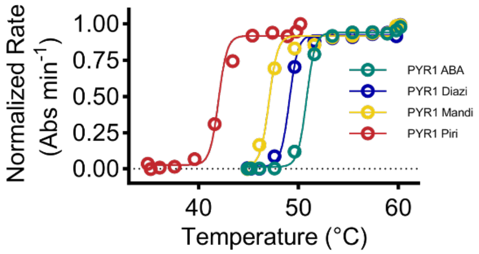


**Supplemental Figure 1 | Thermal stability of engineered PYR1 biosensors.** Phosphatase inhibition of HAB1 as a function of temperature incubation of PYR1. Experiments were performed with 10 µM of ligand (ABA: abscisic acid; Diazi: diazinon; Mandi: mandipropamid; Piri: pirimiphos ethyl), and 50 nM each of the 6xHis-MBP-ΔN-HAB1-C186S-C274S and the indicated engineered PYR1 biosensor. PYR1 designs were incubated for 20 minutes at the indicated temperature. Data is presented as the mean of three technical replicates. Rate is normalized to the maximum rate in the absence of PYR1 protein.

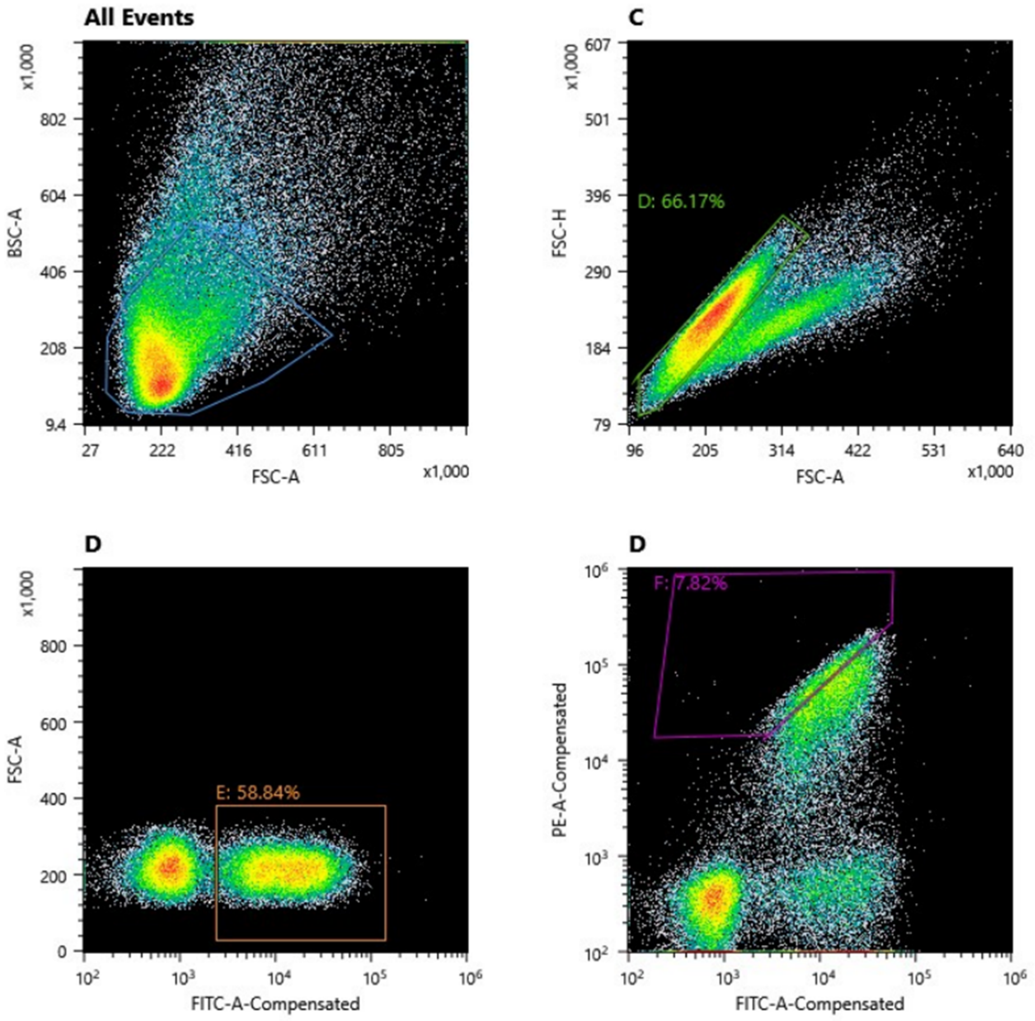


**Supplemental Figure 2 | Representative cell sorting gates used for sorting of PYR1 libraries.** Cells were first gated for yeast cells (top left), single cells (top right), and cells displaying PYR1 variants (bottom left) before sorting for high HAB1-binding cells (bottom right).

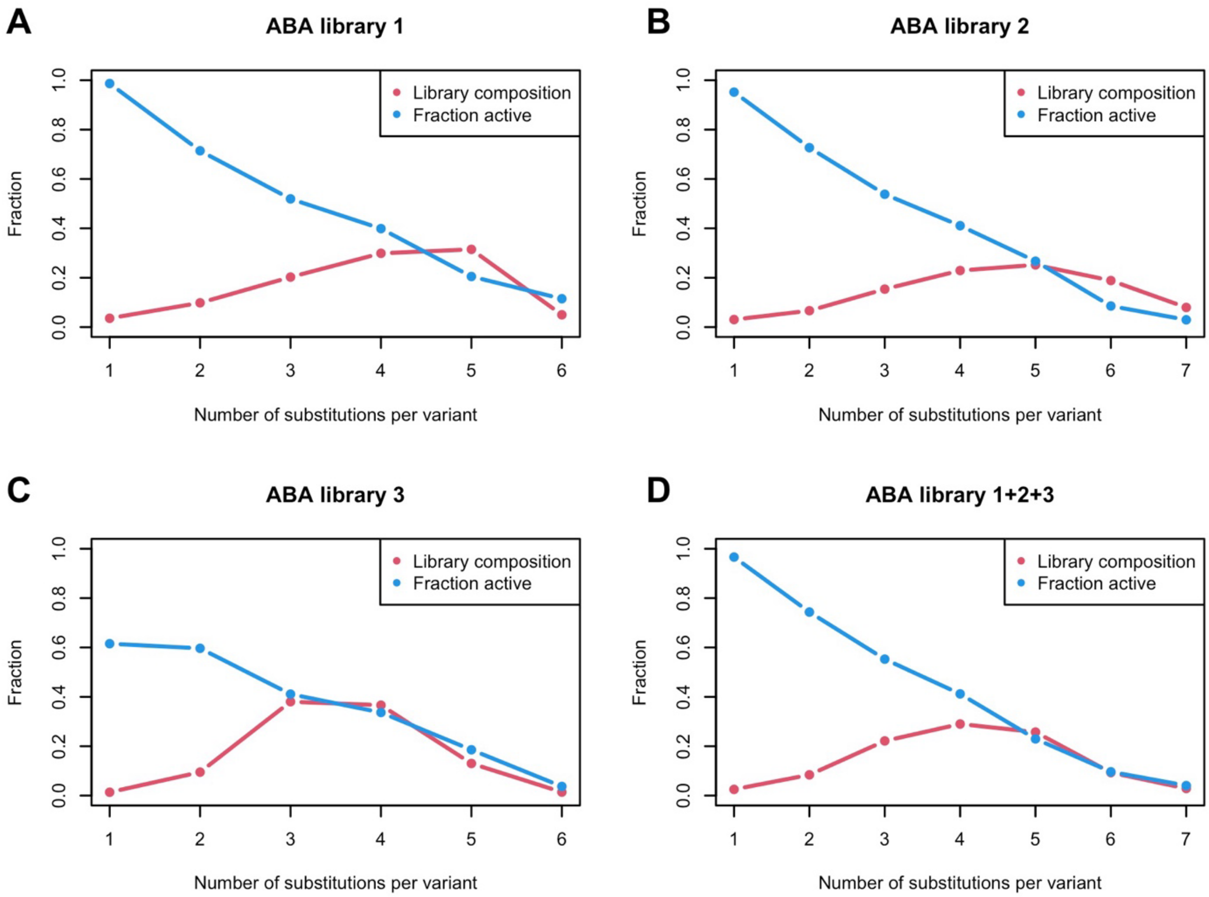


**Supplemental Figure 3 |** Multi-mutant composition of PYR1$^{ABA}$ libraries (red) and the fraction of binding variants (blue) as a function of the number of substitutions per variant. All data sets are appropriate for GMMA analysis with decreasing activity (the average substitution is slightly destabilizing) and a multi-mutant composition that spans the transition from active to largely inactive.

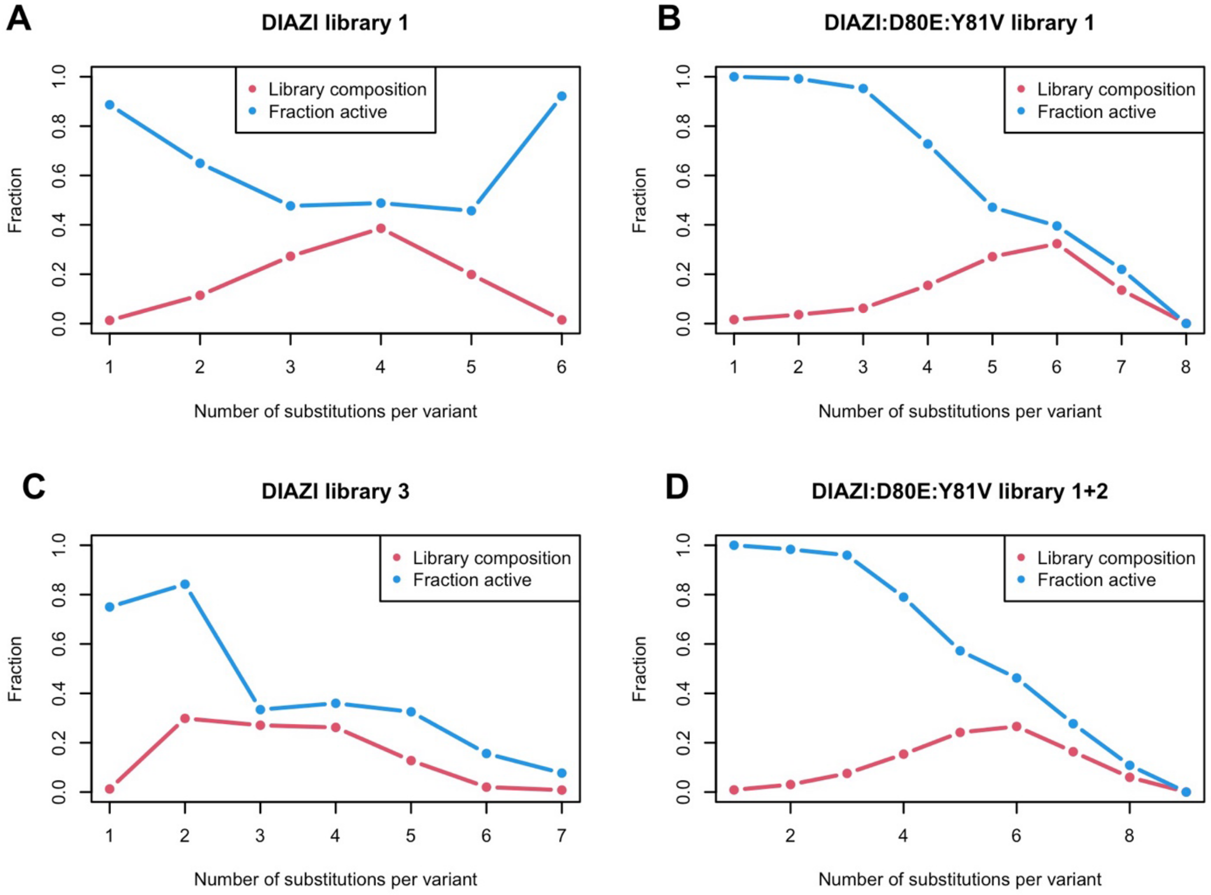


**Supplemental Figure 4 |** Multi-mutant composition of PYR1$^{DIAZI}$ libraries (red) and the fraction of binding variants (blue) as a function of the number of substitutions per variant.

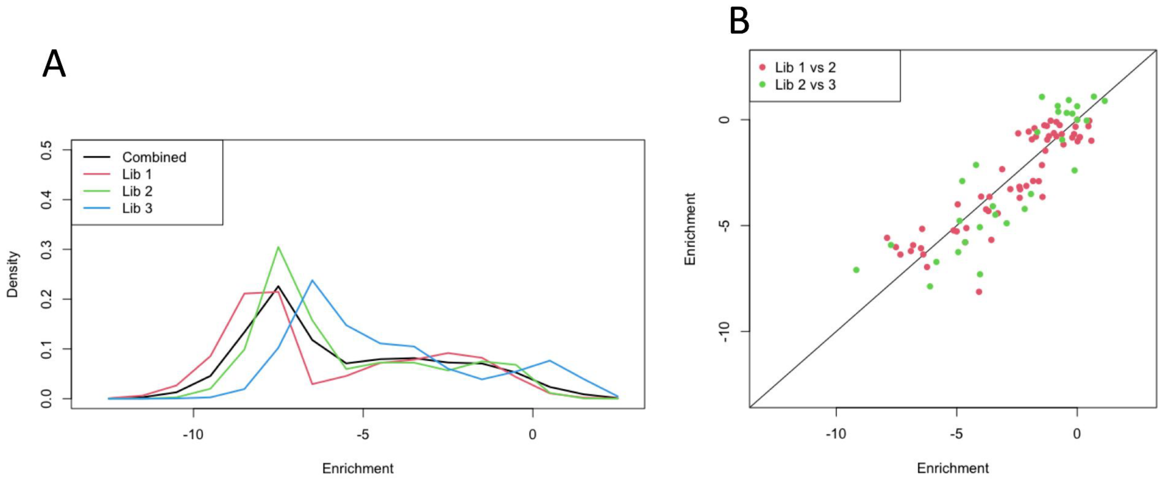


**Supplemental Figure 5 | Enrichment scores of PYR1$^{ABA}$ libraries**. (A) Score distributions of enrichment scores are similar for the three libraries. (B) Enrichment scores of variants that are present in two overlapping libraries are reproduced equally well across the scale, demonstrating the reproducibility of the assay.

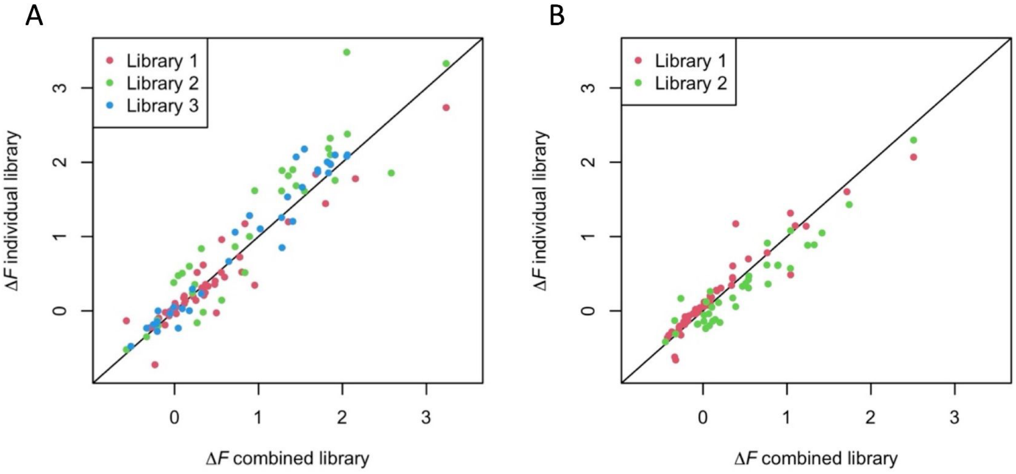


**Supplemental Figure 6 |** Correlation between GMMA effects estimated from individual tile libraries or the combined set of enrichment scores for (A) PYR1$^{ABA}$ and (B) PYR1$^{DIAZI/D80E/Y81V}$. Substitution effects are reproduced well, in particular for the stabilizing substitutions. For PYR1$^{ABA}$, two notable exceptions from library 1 are S47D which is rank 1 in library 1 but rank 8 in the combined library, and D80E which is rank 1 in the combined library but rank 8 in the library 1.

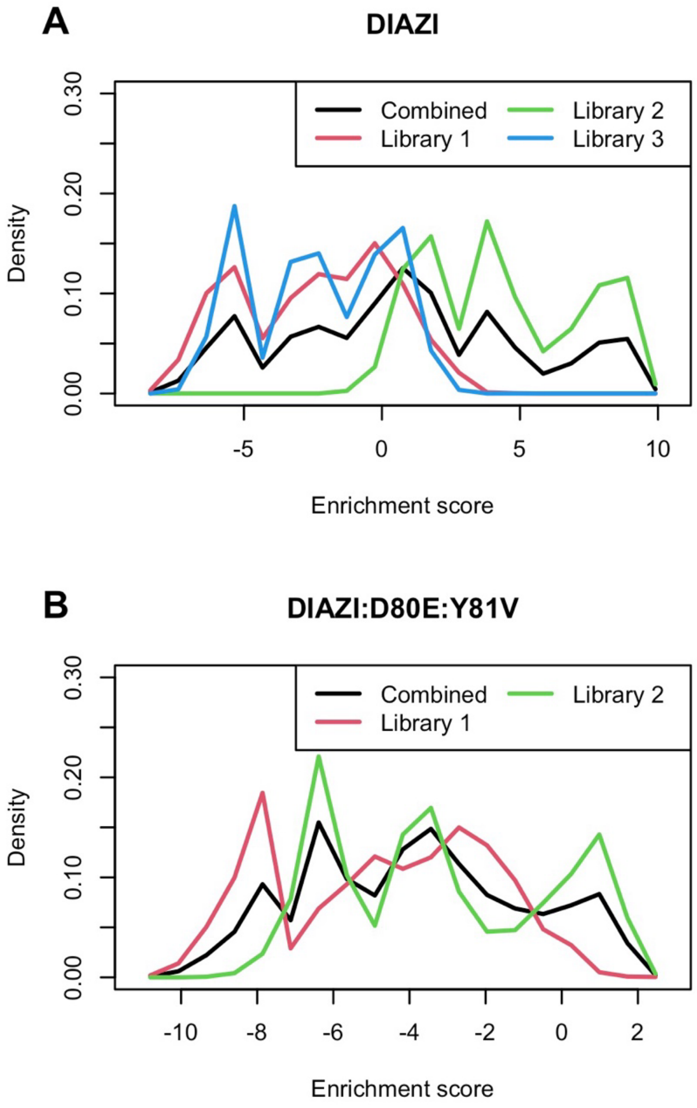


**Supplemental Figure 7 |** Enrichment scores of Diazi sensor libraries. (A) Score distributions are similar for libraries 1 and 3 whereas library 2 has the majority of the library enriched compared to wild type. (B) The substitutions D80E and Y81V are present in both library 1 and 2 and may be used as in an alternative reference wild type for normalization resulting in a score distribution where most variants perform worse than wildtype (score zero). As no variants in library 3 contain Y81V or D80E, library 3 cannot be alternatively referenced.

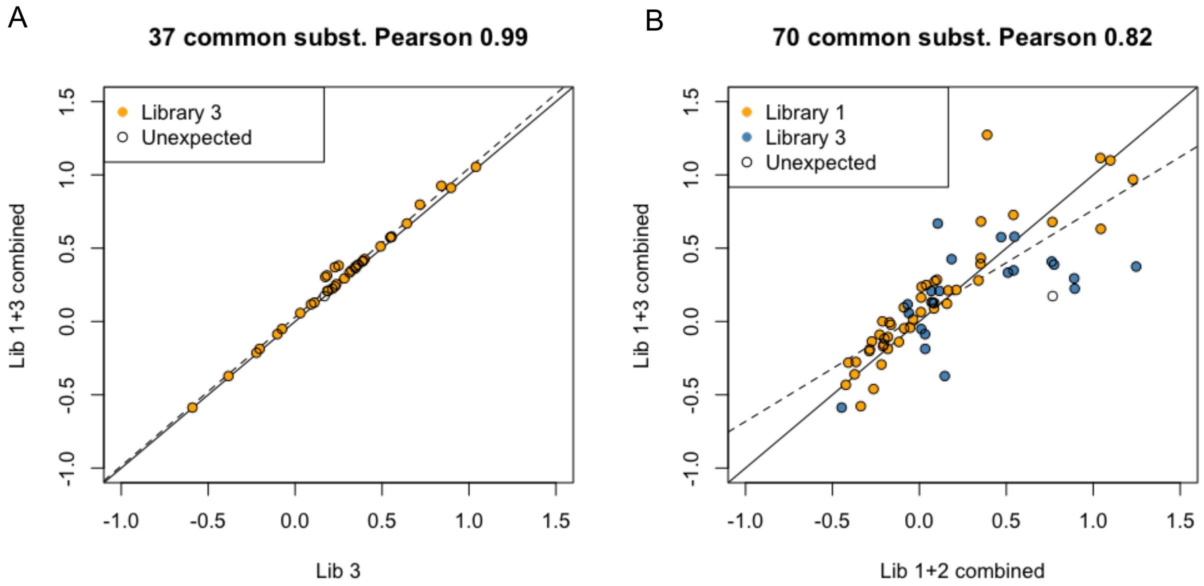


**Supplemental Figure 8 |** GMMA results of PYR1$^{DIAZI}$ tile libraries. (A) Comparison of 37 common substitution effects as determined from library 3 alone or from a combined analysis of library 1+3. The perfect correlation is a result of the disconnected libraries that does not have any substitutions in common (except for four unexpected and rare substitutions that are slightly affected). This indicates that nothing is gained from combining these. (B) Comparison of 70 common substitution effects as determined from a combined analysis of libraries 1+2 and 1+3. The differences arise because the additional substitutions in library 2 are coupled to those of library 1. However, the effects are reproduced well with a Pearson correlation of 0.83.

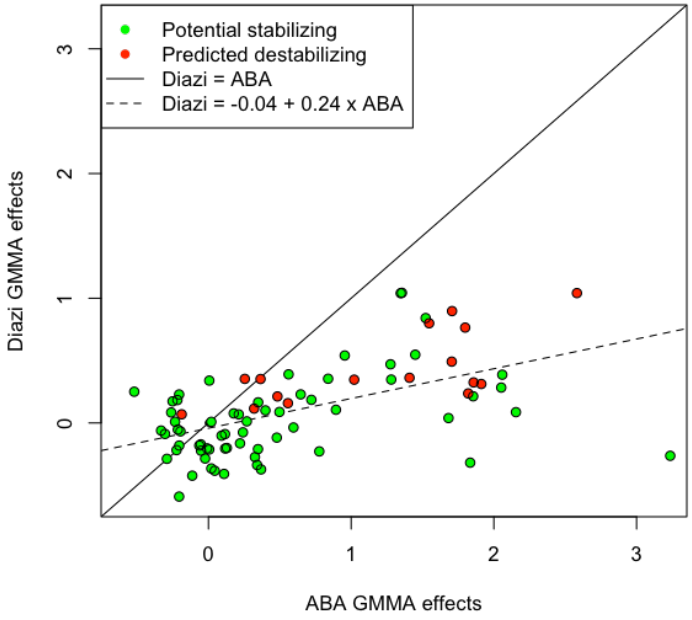


**Supplemental Figure 9 |** GMMA effect from the combined analysis showing all substitutions with confident effects. A line fit with estimated intercept of -0.04 indicates that the zero-point is reproduced well between the two sensors but that effects are on a different scale. A line-fit through zero that ABA effects are 4.7 times higher (not shown).

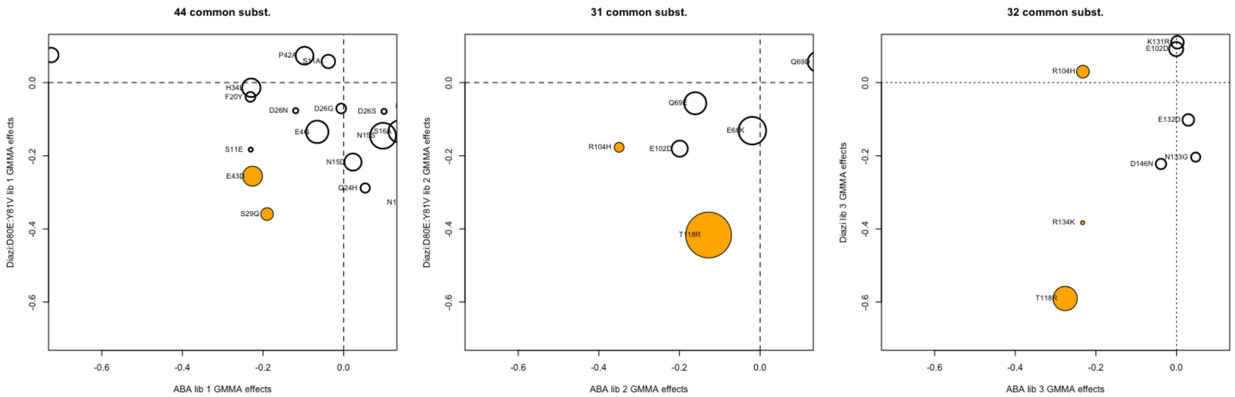


**Supplemental Figure 10 |** Confident GMMA effects for the two sensors compared per tile library. Selected substitutions are highlighted in yellow.

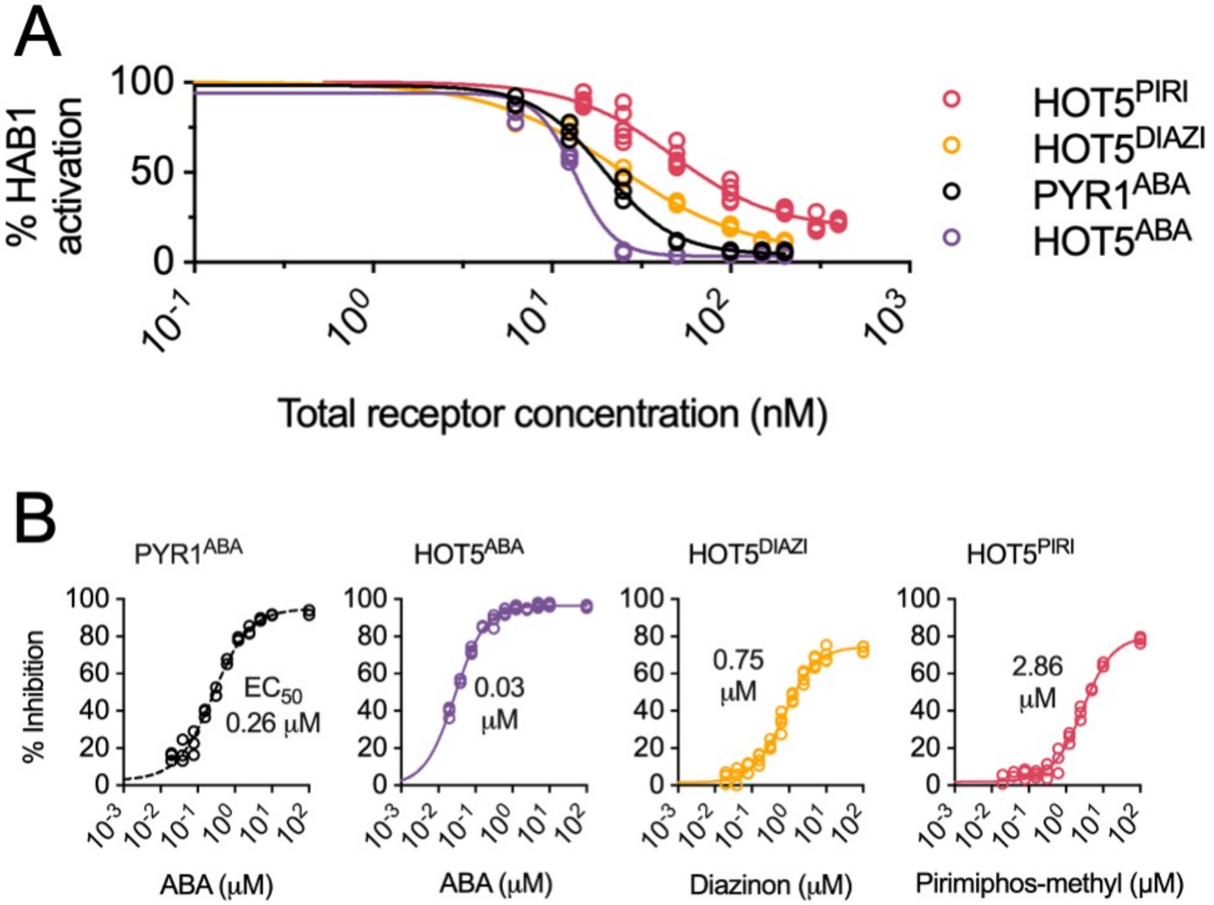


**Supplemental Figure 11 | A.** Protein titration to determine the relative concentration of active receptor to phosphatase protein. Phosphatase inhibition assays were conducted in the presence of ΔN-HAB1 (50 nM for $PYR1^{WT}$, $HOT5^{WT}$, $HOT5^{DIAZI}$, 20 nM for $HOT5^{PIRI}$), 4-methylumbelliferone phosphate substrate (1 mM), varying concentrations of MBP-tagged recombinant receptor proteins (0 nM to 200 nM, or 0 nM to 400 nM for $PYR1^{PIRI}$), and their corresponding ligands (10 uM), ABA, diazinon, or pirimiphos methyl. Curves were fitted to a 4-parameter log-logistic model in Prism GraphPad. Corresponding $EC_{50}$ values of recombinant receptors were used to determine relative ratio of active receptor to phosphatase. **B.** Phosphatase inhibition assays using wild type and HOT5 receptors. Ligand-dependent inhibition of phosphatase activity was assessed using MBP-tagged recombinant receptors (20 nM), ΔN-HAB1 (10 nM), the fluorogenic phosphatase substrate 4-methyl umbelliferone phosphate (MUP; 1 mM), and varying concentrations of target ligands. Inhibition is expressed relative to mock controls ($n = 4$). The curves and inset $EC_{50}$ values are from fits of the dose-response data to a 4-parameter log-logistic model in Prism GraphPad.

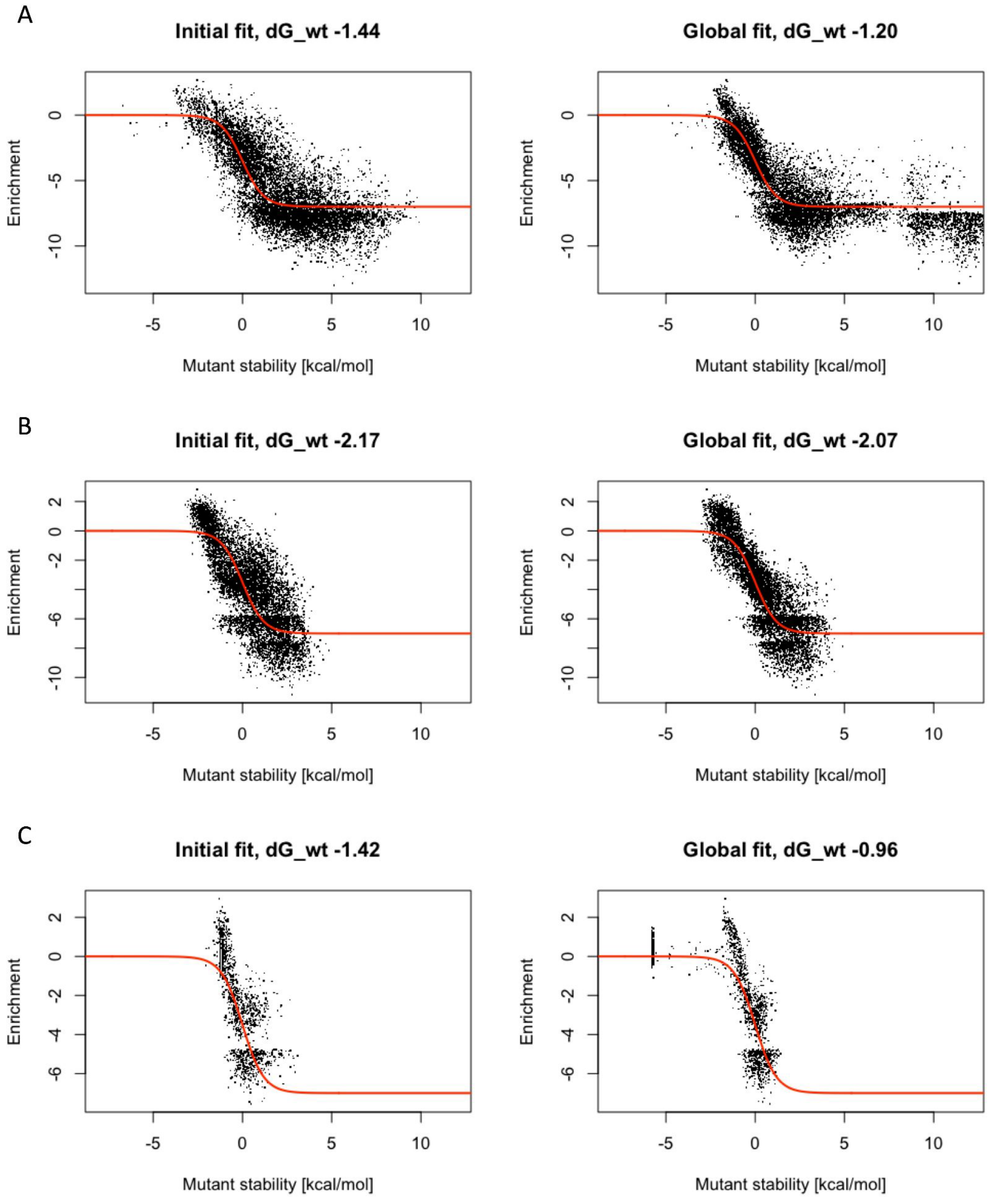


**Supplemental Figure 12 |** GMMA initial and global fit of (A) 9324 variants of PYR1$^{ABA}$ combined libraries 1+2+3, (B) 7839 variants of PYR1$^{DIAZI/D80E/Y81V}$ libraries 1+2, and (C) 1548 variants of PYR1$^{DIAZI}$ library 3.

#### Supplemental Tables

#### Supplemental Table 1. List of mutations used in GMMA.

| Position | WT | MUT | Purpose | Source |
|---|---|---|---|---|
| 4 | E | G | Potential stabilizing | PROSS |
| 9 | E | S | Predicted destabilizing | |
| 11 | S | E/A | Potential stabilizing | PROSS |
| 13 | L | A | Predicted destabilizing | |
| 14 | K | E/A | Potential stabilizing | PROSS |
| 15 | N | D/E/S/Q | Potential stabilizing | PROSS |
| 16 | S | A/F | Potential stabilizing | PROSS |
| 17 | I | V | Potential stabilizing | Highly conserved |
| 18 | A | K/T | Potential stabilizing | PROSS |
| 20 | F | H/Y | Potential stabilizing | PROSS |
| 21 | H | A | Predicted destabilizing | |
| 24 | Q | E/H | Potential stabilizing | PROSS |
| 26 | D | G/N/S/K | Potential stabilizing | PROSS |
| 29 | S | Q/T | Potential stabilizing | PROSS |
| 30 | C | A | Predicted destabilizing | |
| 31 | S | Q | Potential stabilizing | Increase hydrophobic packing |
| 34 | H | L | Potential stabilizing | PROSS |
| 34 | H | A | Predicted destabilizing | |
| 35 | A | I | Potential stabilizing | PROSS |
| 38 | I | A | Predicted destabilizing | |
| 39 | H | N | Potential stabilizing | PROSS |
| 42 | P | A | Predicted destabilizing | |
| 43 | E | D | Potential stabilizing | PROSS |
| 45 | V | A | Predicted destabilizing | |
| 47 | S | D | Potential stabilizing | Increase external charge/polarity |
| 48 | I | V | Potential stabilizing | Highly conserved |
| 48 | I | A | Predicted destabilizing | |
| 52 | F | V | Predicted destabilizing | |
| 54 | K | N | Potential stabilizing | PROSS |
| 68 | E | K | Potential stabilizing | PROSS |
| 69 | Q | E/D | Potential stabilizing | PROSS |
| 70 | N | G | Potential stabilizing | PROSS |
| 71 | F | V | Predicted destabilizing | |

| | | | | |
|---|---|---|---|---|
| 73 | M | Q | Potential stabilizing | Increase external charge/polarity |
| 75 | V | T/E | Potential stabilizing | Increase external charge/polarity |
| 80 | D | E | Potential stabilizing | PROSS |
| 81 | V | Y | DIAZI reversion | |
| 82 | I | N | Potential stabilizing | Highly conserved |
| 87 | L | M | DIAZI reversion | |
| 97 | D | E | Potential stabilizing | PROSS |
| 98 | I | M | Potential stabilizing | Increase external charge/polarity |
| 100 | D | N | Potential stabilizing | Reduced buried charge |
| 102 | E | D | Potential stabilizing | PROSS |
| 102 | E | D | DIAZI reversion | |
| 104 | R | H | Potential stabilizing | PROSS |
| 105 | V | A | Predicted destabilizing | |
| 106 | T | V/I | Potential stabilizing | Increase hydrophobic packing |
| 108 | F | Y | DIAZI reversion | |
| 111 | I | M/E | Potential stabilizing | Increase external charge/polarity |
| 118 | T | R | Potential stabilizing | Highly conserved |
| 121 | K | R/H | Potential stabilizing | PROSS |
| 123 | V | L/F | Potential stabilizing | Increase hydrophobic packing |
| 123 | V | A | Predicted destabilizing | |
| 124 | T | A | Predicted destabilizing | |
| 126 | V | A | Predicted destabilizing | |
| 128 | R | E | Potential stabilizing | PROSS |
| 129 | F | V | Predicted destabilizing | |
| 131 | K | R | Potential stabilizing | Highly conserved |
| 132 | E | D | Potential stabilizing | PROSS |
| 133 | N | G | Potential stabilizing | PROSS |
| 134 | R | K | Potential stabilizing | PROSS |
| 136 | W | H | Predicted destabilizing | |
| 140 | L | A | Predicted destabilizing | |
| 141 | E | S | Predicted destabilizing | |
| 145 | V | A | Predicted destabilizing | |
| 146 | D | N | Potential stabilizing | Reduced buried charge |
| 147 | M | H | Potential stabilizing | Increase external charge/polarity |
| 153 | E | Q | Potential stabilizing | Reduced buried charge |
| 156 | T | A | Predicted destabilizing | |

| | | | | |
|---|---|---|---|---|
| 158 | M | V | DIAZI reversion | |
| 158 | M | R | Potential stabilizing | Increase external charge/polarity |
| 159 | F | G | DIAZI reversion | |
| 160 | A | V | DIAZI reversion | |
| 168 | L | A | Predicted destabilizing | |
| 170 | K | M | Potential stabilizing | Reduced buried charge |
| 171 | L | F | Potential stabilizing | Increase hydrophobic packing |
| 171 | L | A | Predicted destabilizing | |
| 176 | E | Q | Potential stabilizing | Reduced buried charge |
| 178 | M | R | Potential stabilizing | Increase external charge/polarity |
| 179 | A | V/T | Potential stabilizing | Increase hydrophobic packing |

**Supplemental Table 2 | Summary of GMMA results.** All stabilizing (green) and discussed mutations for the combined analyses of both sensors. Bolded mutations are selected for experimental testing.

| **PYR1 ABA lib 1+2+3** | | | | | |
|---|---|---|---|---|---|
| Rank | Subst. | ***ΔF*** | *δ* | $N_{obs}$ | ***ΔF***$_{init}$ |
| 1 | **D80E** | -0.57 | 0.08 | 1298 | -0.39 |
| 2 | M178R | -0.52 | 0.12 | 636 | 0.00 |
| 3 | **R104H** | -0.33 | 0.12 | 304 | -1.12 |
| 4 | F20Y | -0.30 | 0.22 | 134 | -0.11 |
| 5 | **E43D** | -0.29 | 0.14 | 377 | 0.34 |
| 6 | D26K | -0.26 | 0.24 | 175 | 0.74 |
| 7 | A179T | -0.25 | 0.12 | 462 | 0.17 |
| 8 | S47D | -0.23 | 0.17 | 302 | -0.15 |
| 9 | S11E | -0.22 | 0.33 | 114 | 0.90 |
| 10 | A179V | -0.22 | 0.12 | 498 | 0.20 |
| 11 | H34L | -0.21 | 0.15 | 610 | 0.39 |
| 12 | E176Q | -0.21 | 0.16 | 220 | 0.19 |
| 13 | **T118R** | -0.21 | 0.07 | 1275 | -0.45 |
| 14 | D26N | -0.20 | 0.26 | 121 | 0.74 |
| 15 | E102D | -0.20 | 0.11 | 350 | -1.03 |
| 16 | P42A | -0.19 | 0.14 | 424 | 0.57 |
| 17 | **S29Q** | -0.11 | 0.18 | 213 | 0.12 |
| 18 | N15E | -0.11 | 0.31 | 31 | -0.28 |
| 19 | E4G | -0.06 | 0.12 | 421 | 0.56 |
| 20 | D26G | -0.05 | 0.21 | 305 | 1.06 |
| 21 | D146N | -0.05 | 0.19 | 128 | -0.86 |
| 22 | N15D | -0.02 | 0.16 | 256 | 0.58 |
| 23 | N133G | -0.01 | 0.10 | 612 | 0.14 |
| 27 | Q24H | 0.02 | 0.22 | 253 | 0.56 |
| 28 | **R134K** | 0.04 | 0.11 | 851 | 0.41 |
| 30 | N15Q | 0.11 | 0.27 | 61 | 0.28 |
| 32 | N15S | 0.12 | 0.13 | 432 | 0.69 |
| 33 | S16A | 0.13 | 0.14 | 331 | 0.65 |
| 36 | K14E | 0.22 | 0.15 | 295 | 0.77 |
| 41 | Q24E | 0.33 | 0.25 | 210 | 0.74 |
| 42 | E68K | 0.34 | 0.08 | 900 | 0.42 |
| 43 | I17V | 0.35 | 0.15 | 200 | 0.85 |
| 46 | A18T | 0.37 | 0.26 | 67 | 0.55 |
| 83 | V75T | 3.24 | 0.53 | 647 | 3.26 |

| **PYR1 Diazi:D80E:Y81V library 1+2** | | | | | |
|---|---|---|---|---|---|
| Rank | Subst. | ***ΔF*** | *δ* | $N_{obs}$ | ***ΔF***$_{init}$ |
| 1 | **T118R** | -0.45 | 0.07 | 1317 | -0.34 |
| 2 | **S29Q** | -0.42 | 0.15 | 156 | 0.32 |
| 3 | N15Q | -0.41 | 0.32 | 28 | 0.66 |
| 4 | A18T | -0.37 | 0.22 | 49 | 0.21 |
| 5 | Q24H | -0.36 | 0.16 | 181 | 0.74 |
| 6 | E68K | -0.34 | 0.07 | 1194 | -0.16 |
| 7 | E80D | -0.33 | 0.59 | 5995 | 0.70 |
| 8 | **E43D** | -0.29 | 0.13 | 214 | 0.40 |
| 9 | N15D | -0.29 | 0.12 | 229 | 0.51 |
| 10 | Q24E | -0.27 | 0.17 | 160 | 0.74 |
| 11 | V75T | -0.26 | 0.07 | 894 | 0.51 |
| 12 | F20H | -0.23 | 0.26 | 35 | 0.46 |
| 13 | S11E | -0.22 | 0.24 | 59 | 0.59 |
| 14 | D26S | -0.21 | 0.25 | 66 | 0.86 |
| 15 | I17V | -0.21 | 0.12 | 236 | 0.50 |
| 16 | N15S | -0.20 | 0.10 | 398 | 0.56 |
| 17 | S16A | -0.20 | 0.11 | 312 | 0.49 |
| 18 | D26N | -018 | 0.25 | 66 | 1.03 |
| 19 | E4G | -0.18 | 0.12 | 230 | 0.54 |
| 20 | D26G | -0.17 | 0.17 | 178 | 0.84 |
| 21 | K14E | -0.16 | 0.15 | 131 | 0.68 |
| 24 | F20Y | -0.09 | 0.15 | 123 | 0.50 |
| 25 | E102D | -0.07 | 0.11 | 419 | -0.40 |
| 26 | **R104H** | -0.06 | 0.16 | 223 | 0.26 |
| 27 | H34L | -0.05 | 0.11 | 475 | 0.59 |
| 30 | S47D | 0.01 | 0.12 | 239 | 0.46 |
| 37 | P42A | 0.07 | 0.13 | 261 | 0.57 |
| **PYR1 Diazi library 3** | | | | | |
| 1 | **T118R** | -0.59 | 0.12 | 281 | -0.19 |
| 2 | **R134K** | -0.38 | 0.20 | 77 | 0.31 |
| 3 | D146N | -0.22 | 0.14 | 212 | -0.13 |
| 4 | N133G | -0.20 | 0.16 | 128 | 0.21 |
| 7 | **R104H** | 0.03 | 0.19 | 79 | 0.12 |
| 11 | A179T | 0.17 | 0.22 | 462 | 0.19 |
| 16 | E176Q | 0.23 | 0.25 | 83 | 0.30 |
| 19 | M178R | 0.25 | 0.22 | 538 | 0.31 |

#### Supplemental Table 3: Strains used in the study

| Plasmid | Description | Source |
|---|---|---|
| *E. coli* TOP 10 | F- *mcrA* Δ (*mrr-hsd*RMS-*mcr*BC) Φ80*lac*ZΔM15 Δ *lac*X74 *rec*A1 *ara*D139 Δ (araleu)7697 *gal*U *gal*K *rps*L (StrR) *end*A1 *nup*G | Thermo Fisher Scientific |
| YS626 | *K. marxianus* CBS 6556 *ura3Δ his3Δ* | Löbs et al. |
| YS2042 | YS626 *abz1*::Z4BS-HTB1core-*GFP-CYC1*t | This study |
